# Molecular Glue-like Degraders of TEM β-Lactamases by Periplasmic Protease DegP

**DOI:** 10.64898/2026.03.30.715243

**Authors:** Emilia K. Taylor, Patricia Santos Barbosa, Tarini Kadambi, Frederik Eisele, Elisabete C. C. M. Moura, Timothy R. Walsh, Georgia L. Isom, Thomas Lanyon-Hogg

## Abstract

Antimicrobial resistance is one of the most serious challenges to global health, yet the development of new molecules with novel mechanisms of action to combat resistance is lacking. Here, we report the discovery of molecular glue-like compounds that recruit TEM-family β-lactamases to the bacterial protease DegP for degradation. β-lactamase inhibitor tazobactam was found to accelerate degradation of TEM β-lactamases by DegP, which was further enhanced by linkerless incorporation of dipeptide motifs enriched among DegP substrates. The resulting molecular glue-like degraders showed improved synergy with β-lactam piperacillin against resistant *E. coli* compared to tazobactam, as well as good pharmacokinetic properties for oral dosing. Collectively, this work establishes periplasmic targeted protein degradation as a promising new mechanism for combating β-lactamase resistance.

## Introduction

Antimicrobial resistance (AMR) is a critical threat to health globally, which undermines both antibiotic effectiveness and many common medical procedures.[1] Many antibiotics currently in development are derivatives of known antibiotic classes and are therefore rapidly compromised by existing resistance mechanisms, meaning there is an urgent need for new molecules functioning through novel new mechanisms of action to combat resistance. This includes development of both direct-acting antibiotics, as well as antibiotic adjuvants that may slow or reverse the evolution of resistance.[2]

Targeted protein degradation (TPD) is an emerging strategy to harness endogenous cellular proteolytic systems to eliminate disease-causing proteins.[3] Bifunctional proteolysis-targeting chimera (PROTAC) degraders have demonstrated utility in eukaryotic systems, but the application of TPD in bacteria has remained limited to cytoplasmic proteases (ClpXP, ClpCP) and largely depends on engineered substrates.[4–7] PROTACs offer several potential mechanistic advantages compared to small-molecule inhibitors: proteolytic removal of target proteins may lead to longer phenotypic effects; degraders may eliminate multiple copies of the target protein leading to a catalytic effect; binding sites not suited to small molecule inhibition may be amenable to PROTAC targeting as degradation can be achieved with lower-affinity, transient binders of shallow pockets or allosteric sites. However, the large size of PROTAC molecules (typically >1,000 Da) present challenges in pharmacokinetics, and has led to increasing interest in the development of molecular glues. Molecular glues are monovalent small molecules that induce conformational changes to promote new target interactions or stabilise weak existing interactions; however, the discovery of molecular glues has historically been serendipitous.[8]

β-lactamase enzymes hydrolyse β-lactam antibiotics (Figure 1A), representing one of the most prevalent antibiotic resistance mechanisms in Gram-negative bacteria, with >2,000 enzyme variants identified to-date.[9] Covalent inhibitors of serine β-lactamase are used clinically; however the efficacy of these compounds can be compromised by β-lactamase mutation or overexpression.[10] A recent study developed a PROTAC that recruited the cytoplasmic protease ClpXP to degrade the CTX-M-14 β-lactamase, which had been modified to remain in the cytoplasm. However, as β-lactamases are located in the periplasm of Gram-negative bacteria, ClpXP-mediated degradation is not applicable to native β-lactamases.[5] Further, the resulting PROTAC incorporates the ClpXP-recruiting SsrA tag (AANDENYALAA) into β-lactamase inhibitor nacubactam, and is therefore high-molecular-weight (1,669 Da). There is therefore an unmet need for degrader molecules that target β-lactamase resistance enzymes in their native periplasmic compartment, and which possess low molecular weight and favourable physicochemical properties.

**Figure 1.**
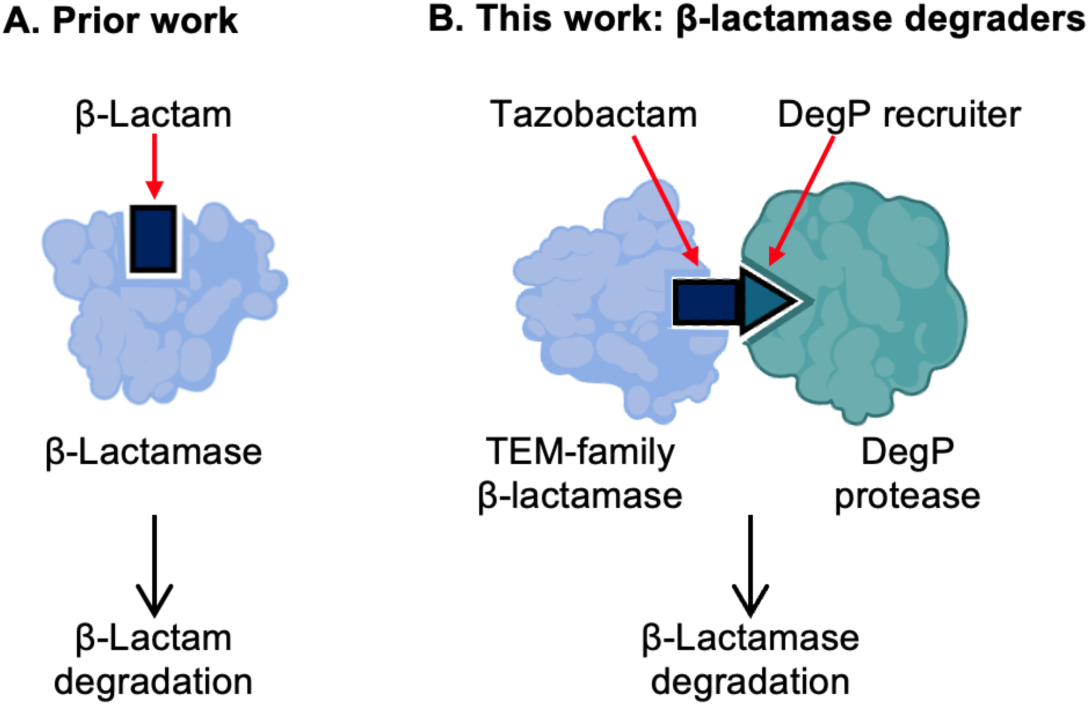
Schematic overview of targeted protein degradation strategy in *E. coli*. A) β-Lactam antibiotics, and at a slower rate covalent β-lactamase inhibitors, are degraded by β-lactamase enzymes. B) Proposed novel mechanism of action developed in this work, by converting tazobactam into molecular glue-like degraders that recruit TEM-family β-lactamases to the DegP protease for degradation.

DegP is a serine protease required for the maintenance of proteostasis in the periplasm of Gram-negative bacteria, primarily targeting misfolded and aggregated proteins.[11] In *E. coli,* DegP forms a self-assembling, substrate-activated proteolytic cage that recognises hydrophobic patches and C-terminal residues of target proteins.[12] To date, DegP, and more broadly the periplasm, has not been explored for PROTAC development. Here, we report the first molecular glue-like degraders of β-lactamases via recruitment of periplasmic protease DegP (Figure 1B). These molecules resensitise resistant bacteria and possess suitable pharmacokinetic properties for oral dosing.

## Results

Initially, the ability of DegP to degrade three clinically-relevant β-lactamases *in vitro* was assessed, without additional degrader molecules. Purified serine β-lactamases TEM-116 (class A), AmpC (class C), and OXA-48 (class D), were incubated with purified DegP for 24 h and analysed by SDS-PAGE. TEM-116 was almost completely degraded within 24 h, whereas OXA-48 and AmpC remained intact (Figure 2A). TEM-1, which differs from TEM-116 by a single point mutation, was also degraded by DegP (Figure S1A).[13] To confirm that degradation was dependent on DegP catalytic activity, TEM-116 was incubated with either catalytically-inactive S210A DegP or DegP pre-treated with the serine protease inhibitor diisopropyl fluorophosphate. TEM-116 degradation was blocked in both conditions, confirming that DegP activity was required for loss of TEM-116 (Figure 2B). A decrease in DegP was also observed in the presence of TEM-116, which was attributed to hyperactivation and autocleavage of DegP triggered by TEM-116 degradation products (Figure S1B). As DegP primarily degrades misfolded and aggregated proteins,[11] the structural integrity of TEM-116 was evaluated using differential scanning calorimetry and size-exclusion chromatography (SEC), (Figure S1C-D). This confirmed TEM-116 was folded and not aggregated, and TEM-116 β-lactamase activity was further demonstrated in a nitrocefin hydrolysis assay (Figure S1E).[14] Taken together, these findings showed that DegP can degrade folded and active TEM-116 *in vitro*, suggesting a suitable periplasmic target and protease pair for degrader development. This may further imply that DegP has a broader role in periplasmic protein degradation than previously appreciated, and make DegP an attractive tool for TPD of periplasmic proteins.[15–18]

**Figure 2.**
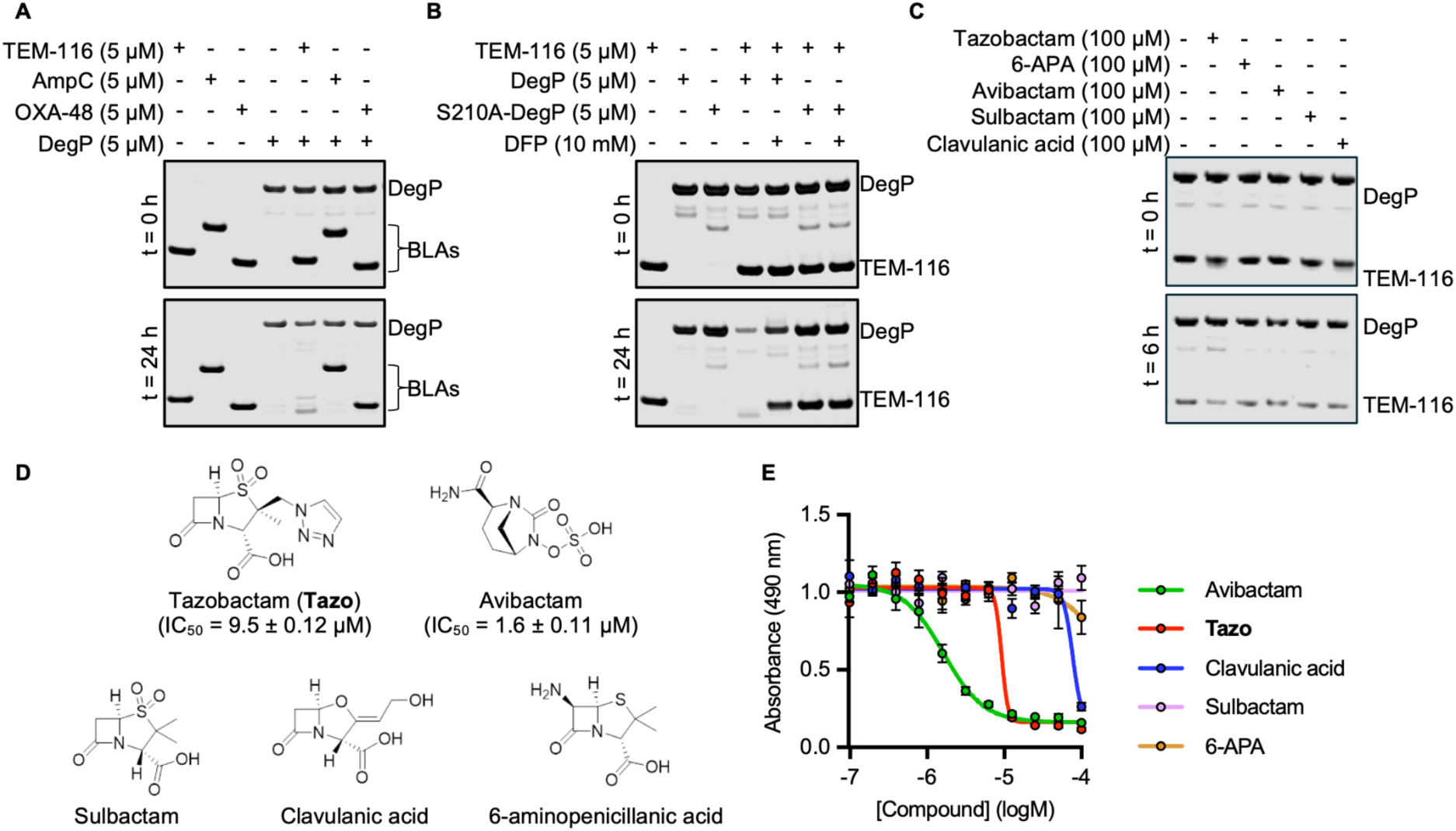
DegP-mediated degradation of TEM-116 β-lactamase. (A) Co-incubation of DegP with β-lactamase *in vitro* for 24 h resulted in a loss of TEM-116, whereas AmpC and OXA-48 β-lactamases were unaffected. (B) TEM-116 degradation required DegP catalytic activity, as the S210A-DegP mutant showed no degradation and protease inhibitor DFP protected TEM-116 from degradation. (C) β-Lactamase inhibitor tazobactam accelerated DegP-mediated degradation of TEM-116 over 6 h co-incubation, compared to other β-lactamase inhibitors or β-lactam precursor 6-aminopenicillanic acid (6-APA). (D) Structures of tested β-lactamase inhibitors and 6-APA, with corresponding IC_50_ values for TEM-116 inhibition. (E) Dose-response analysis of β-lactamase inhibitors and 6-APA in TEM-116 nitrocefin assay. Data represent mean ± SEM (n = 3).

We next sought to design molecules to enhance degradation of TEM by DegP. To identify a suitable ligand for TEM-116, a panel of four clinical β-lactamase inhibitors (tazobactam, avibactam, sulbactam, and clavulanic acid) and a precursor in β-lactam synthesis (6-aminopenicillanic acid (6-APA)) were screened to evaluate their effect on DegP-mediated degradation of TEM-116 over 6 h. Clavulanic acid, sulbactam, 6-APA, and avibactam did not affect degradation; however, surprisingly, tazobactam enhanced TEM-116 degradation (Figure 2C). Both avibactam and tazobactam showed dose-dependent inhibition of TEM-116 in the nitrocefin hydrolysis assay (IC_50_ = 1.6 ± 0.11 µM and 9.5 ± 0.12 µM, respectively), whereas sulbactam, 6-APA, and clavulanic acid showed no inhibition <100 µM (Figure 2D,E). None of the β-lactamase inhibitors induced degradation of OXA-48 or AmpC by DegP (Figure S2A), although higher concentrations of tazobactam were required for inhibition (OXA-48 IC_50_ = 41 ± 0.009 µM and AmpC IC_50_ = 52 ± 0.06 µM, Figure S2B,C). To understand the mechanism by which tazobactam accelerated TEM-116 degradation, the integrity of TEM-116 in the presence and absence of tazobactam under degradation assay conditions was assessed by SEC (Figure S1D). Treatment with DMSO alone resulted in the majority of TEM-116 eluting as a single, monodisperse peak after 4 h, representing folded protein. In contrast, tazobactam treatment altered the integrity of TEM-116, with multiple peaks observed by SEC (Figure S1D). This suggested that tazobactam may induce TEM-116 to adopt a more DegP-sensitive conformation, rather than leading to stabilization of TEM-116 as previously reported.[19] Collectively, these data suggested tazobactam may act as a molecular glue to enhance DegP degradation of TEM-116, therefore representing a suitable target-recruiting ligand for further degrader development.

With our unexpected observation that DegP can degrade folded and functional TEM-116, we next sought to identify other proteins that may be susceptible to DegP-mediated degradation to guide degrader development. *E. coli* lysates were incubated with either WT DegP or catalytically-inactive S210A-DegP, followed by tryptic digest and LC-MS/MS analysis. 447 proteins were statistically significantly depleted (log_2_ fold-change <−1, p <0.05) following treatment with WT DegP (Figure 3A). S210A-DegP treatment increased the abundance of 180 proteins (log_2_ fold-change >1, p <0.05, Figure 3B), potentially due to S210A-DegP competition with endogenous DegP for substrate binding. The proteins depleted by addition of WT DegP may represent weakly-engaged substrates that can be degraded at higher local DegP concentrations. DegP recognises the C-terminal of target proteins, and analysis of the ten C-terminal residues of proteins depleted by WT DegP revealed increased prevalence of hydrophobic amino acids at the final two positions: phenylalanine at the penultimate position (P9) and alanine, leucine, or isoleucine at the terminal position (P10) (Figure 3C). Additionally, upstream positions showed increased prevalence of proline (P2), serine (P3), histidine (P4), and leucine (P5). These sequences suggested potential degron motifs to enhance DegP recruitment to folded proteins, which could be incorporated into degrader molecules.

**Figure 3.**
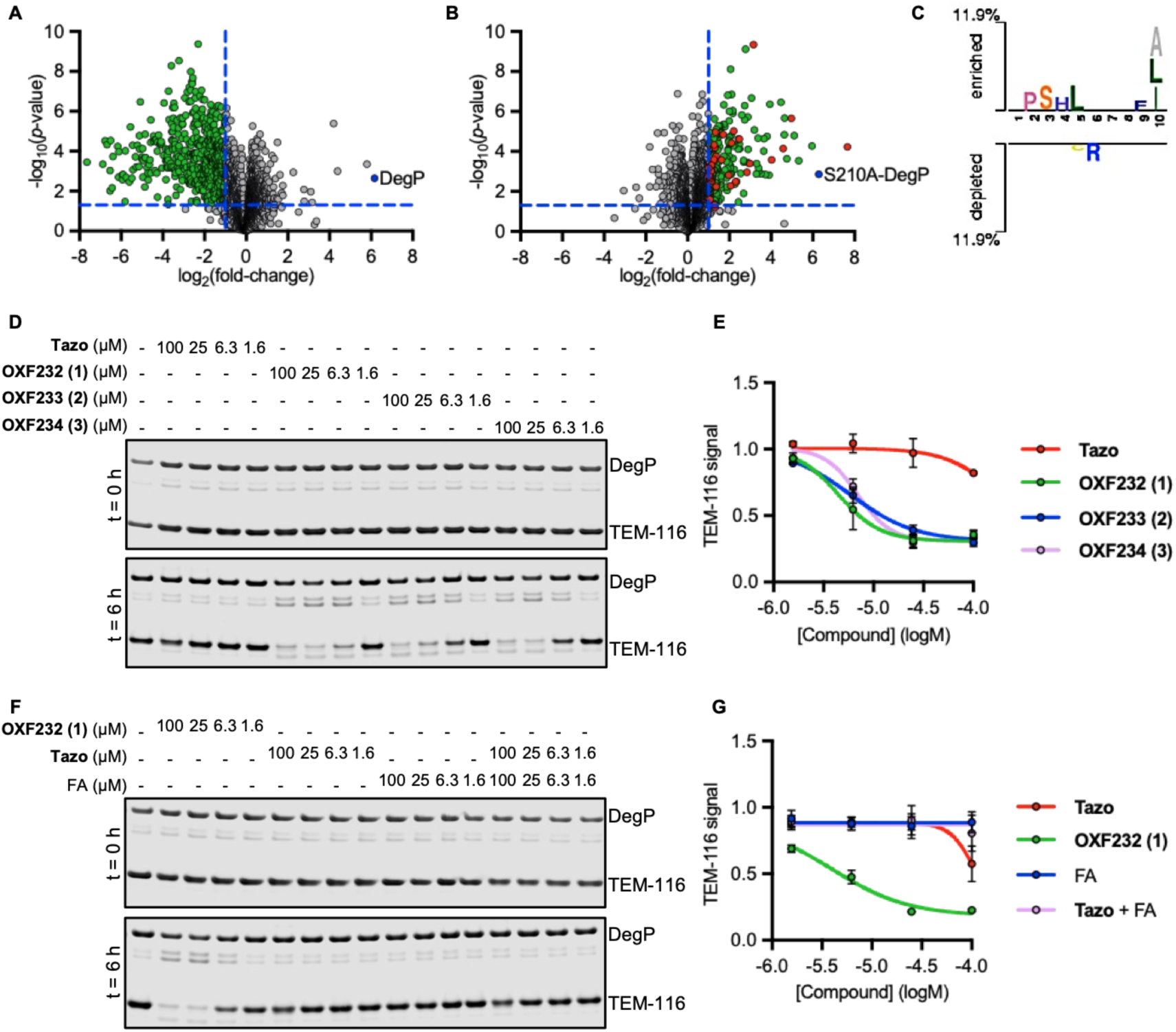
Quantitative global proteomic analysis of DegP substrates and *in vitro* evaluation of optimised degraders. (A) Identification of proteins decreased in abundance in *E. coli* lysate following treatment with WT DegP (log_2_ fold-change <−1, p <0.05). (B) Identification of proteins increased in abundance in *E. coli* lysate following treatment with S210A-DegP (log_2_ fold-change >1, p <0.05). Proteins increased in abundance following both WT and S210A-DegP treatment are shown in red. (C) Sequence analysis of ten C-terminal residues of target proteins identified in (A), highlighting residues with increased or decreased prevelance at each position. (D) Dose-response analysis of TEM-116 degradation over 6 h incubation with triazole-linked degraders. (E) Quantification of TEM-116 degradation dose-response in (D). (F) Dose-response analysis of TEM-116 degradation over 6 h incubation with triazole-linked **OXF232** (**1**) degrader, components tazobactam (**Tazo**), the FA dipeptide, or tazobactam and the FA dipeptide in combination. (G) Quantification of TEM-116 degradation dose-response in (F). Data represent mean ± SEM (n = 3).

Having identified tazobactam as a DegP-TEM-116 molecular glue, and C-terminal dipeptides FA, FI, and FL as enriched in DegP substrates, hybrid degrader molecules were designed to target TEM-116. Two series of degraders, which differed in the attachment position of the FA/FI/FL dipeptide to tazobactam, were synthesised without linker groups to maintain a low molecular weight and glue-like properties (Scheme 1). In the first series, a novel route to triazole-functionalised tazobactam was established via an azide analogue of tazobactam (Scheme S1) and attachment to alkyne-tagged dipeptides using a copper-catalysed azide-alkyne cycloaddition to yield triazole-linked FA (**OXF232, 1**), FI (**OXF233**, **2**), and FL (**OXF234**, **3**) analogues. In the second series, the dipeptides were assembled and coupled to the tazobactam carboxylate group using microwave-assisted solid-phase peptide synthesis to yield amide-linked FA (**4**), FI (**5**), and FL (**6**) analogues.

**Scheme 1.**
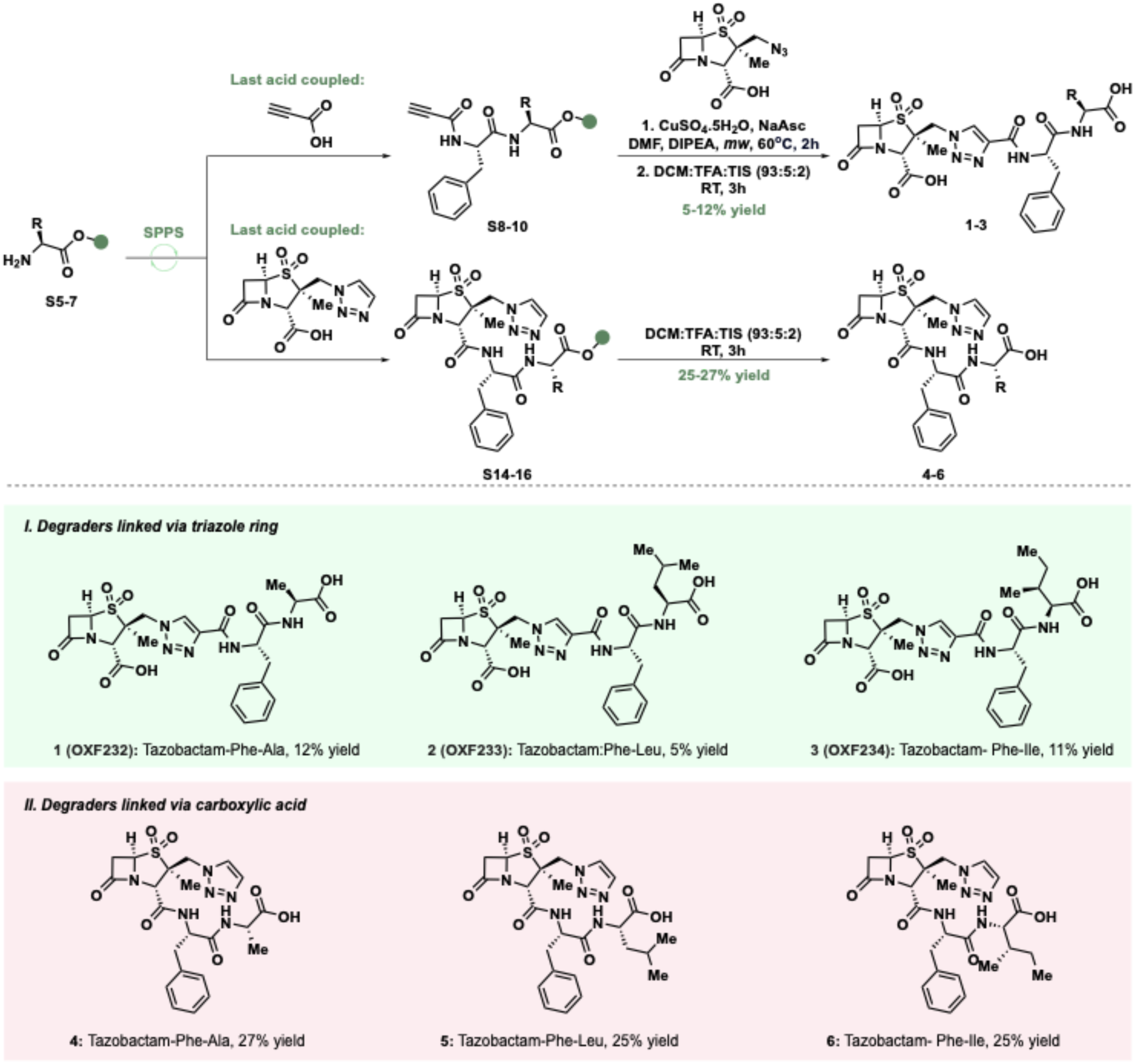
Synthetic route to degrader molecules using solid-phage peptide synthesis

The synthesised compound panel was assessed in dose-response for degrader activity with TEM-116 and DegP over 6 h. Triazole-linked compounds **1**-**3** induced TEM-116 degradation with half-maximal degradation concentration (DC_50_) values of 5.8 ± 0.14 µM (**OXF232**), 6.9 ± 0.13 µM (**OXF233**), and 6.6 ± 0.15 µM (**OXF234**), whereas tazobactam showed DC_50_ >100 µM over this timeframe (Figure 3D). In contrast, the carboxyl-linked compounds **4-6** showed no degradation activity (Figure S3A). TEM-116 inhibition assays demonstrated that triazole-linked degraders **1-3** retained inhibitory activity, whereas amide-linked analogues **4-6** were inactive, consistent with the known essential binding interaction of the carboxylate with β-lactamases (Figure S3B-C).[11] Triazole-linked FA degrader **OXF232** (**1**) was the best performing analogue, and we therefore tested whether enhanced TEM-116 degradation required both components of the degrader to be linked. **OXF232** was compared with tazobactam and the FA dipeptide, either as single agents or in combination. Enhanced degradation was not observed with the unlinked components of **OXF232** over 6 h (Figure 3E), consistent with induced proximity between TEM-116 and DegP, rather than independent binary binding, being required for proteolysis. Overall, these findings identify new dipeptide recruiters of DegP that could be incorporated into other degraders in the future. Furthermore, to our knowledge this is the first demonstration of molecular glue-like molecules being used to enhance degradation of bacterial proteins.

Having demonstrated that the triazole-linked degraders accelerated TEM-116 degradation *in vitro*, the ability of the compounds to sensitise β-lactamase-expressing bacteria to β-lactams was evaluated. Kinetic growth curves were performed in *E. coli* harbouring TEM-1 on a pK18 plasmid, where TEM-1 is expressed from its native constitutive promoter. *E. coli* pK18-TEM-1 exhibited resistance to the β-lactam piperacillin which is commonly co-administered with tazobactam in the clinic, with growth observed at 1 mg/mL piperacillin (Figure S4A). Dose responses of tazobactam or degraders **1-3** were tested in combination with piperacillin (1, 0.33, and 0.1 mg/mL). At lower piperacillin concentrations, tazobactam outperformed the degrader molecules (Figure 4A-B). However, at 1 mg/mL piperacillin, the degraders outperformed tazobactam (Figure 4C) and exhibited improved synergy for bacterial growth inhibition (Table S1). This observation may reflect a mechanism by which increased piperacillin concentrations selects for cells with higher intracellular TEM-1 concentrations. This may favour degrader activity through DegP-mediated elimination of TEM-1, whilst tazobactam is slowly hydrolysed by TEM-1 during inhibition.[20] **OXF232** (**1**), containing the FA dipeptide, outperformed the FI and FL counterparts **OXF233** (**2**) and **OXF234** (**3**), respectively, consistent with *in vitro* degradation assays (Figure 3D). Due to the efficient killing of bacteria under conditions where degraders **1-3** showed enhanced biological activity over tazobactam, it was not possible to obtain sufficient bacteria to test if cellular degradation of TEM-1 had occurred. Taken together, our *in vitro* and *in vivo* data suggest that degraders can sensitise resistant *E. coli* to piperacillin, and offer enhanced potency at high β-lactamase expression levels over the clinically used inhibitor tazobactam.

**Figure 4.**
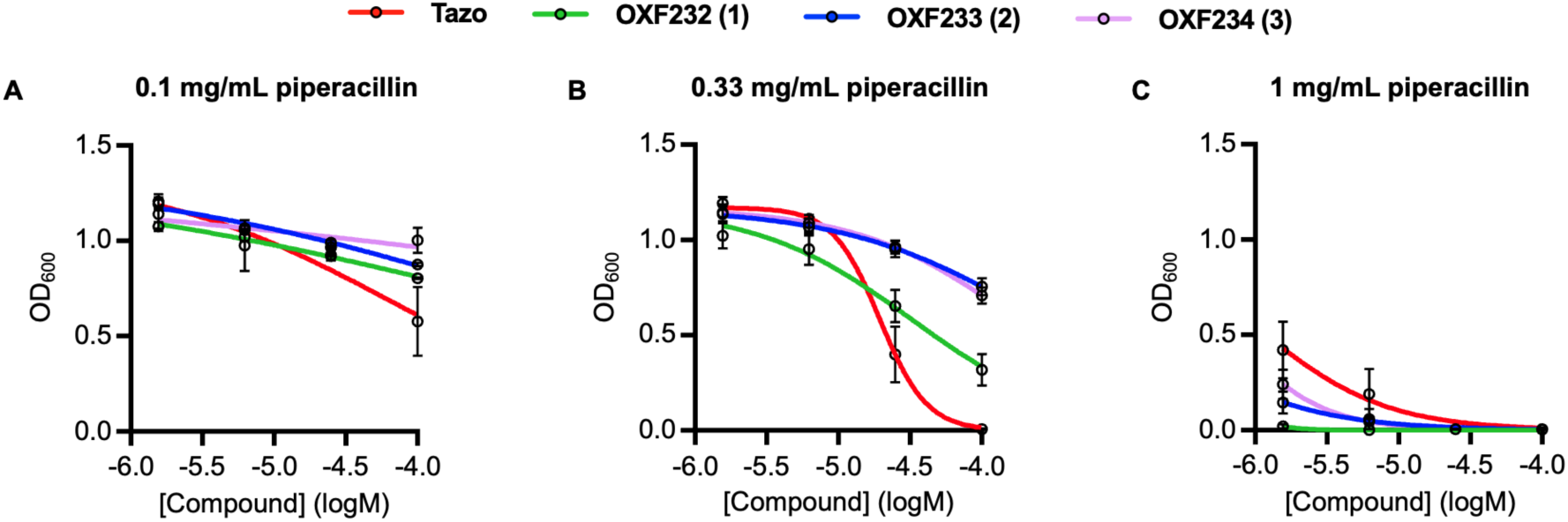
*In vivo* resensitisation of *E. coli* harbouring TEM-1 to piperacillin. Growth of *E. coli* containing pK18-TEM-1 at (A) 0.1 mg/mL piperacillin, (B) 0.33 mg/mL piperacillin, (C) 1 mg/mL piperacillin. Degraders showed improved activity at higher piperacillin levels compared to tazobactam. Data represent mean ± SEM (n = 3).

The large size of traditional PROTAC degraders leads to pharmacokinetic challenges, therefore we assessed if these issues were alleviated by our smaller molecular glue-like degraders. The pharmacokinetic and mammalian cytotoxicity properties of **OXF232** and **OXF233** were evaluated. Both compounds exhibited favourable physicochemical profiles, with high aqueous kinetic solubility (>175 µM in phosphate buffer, pH 7.4) and equilibrium solubility (LogD <-1.4). High metabolic stability was observed in both human and mouse liver microsomes, with calculated half-lives >139 min, suggesting minimal hepatic metabolism (Table S2). Cytotoxicity testing in HEK293T cells demonstrated that **OXF232** and **OXF233** were non-toxic at concentrations <100 µM over 24 h (Figure S4B). Taken together, these data demonstrated that the developed molecular glue-like degraders possessed promising drug-like properties compatible with further pharmacological development, including high aqueous solubility, metabolic stability, and low mammalian toxicity, which are common challenges in PROTAC development (Figure S4B, Table S2).

## Discussion

In summary, we present here the first demonstration of targeted degradation of an antibiotic resistance enzyme in its native cellular location. β-lactamase inhibitor tazobactam was identified as showing molecular glue-like properties by accelerating the degradation of TEM-116 by DegP. The structural basis of this effect, therefore, poses an important question in determining the scope of other β-lactamases or bacterial proteins that may be targeted for degradation. DegP represents an attractive protease for TPD in bacteria, being co-localised with resistance enzymes such as β-lactamases, and further supported by a growing body of evidence for the ability of DegP to degrade folded/functional proteins.[21] Quantitative proteomics identified C-terminal hydrophobic dipeptides as enriched in DegP substrates, which enabled the rational design of low-molecular-weight degraders with favourable physicochemical properties. Alternative protease-recruiter moieties may also lead to enhanced degraders, for example the PSHL motif (Figure 3D) which was not explored in this work, and improved understanding of bacterial protease regulation is therefore an important area for future TPD research. The developed molecular glue-like degraders outperformed clinically-approved β-lactamase inhibitor tazobactam under conditions promoting high β-lactamase expression, suggesting a mechanistic advantage of proteolysis-based degradation compared to covalent inhibitors in such cases.[22] As antimicrobial resistance continues to place an increasing burden on global healthcare, the development of TPD in bacteria may provide a critical new mechanism of action to combat drug-resistant infections.

## Supporting Information

The authors have cited additional references within the Supporting Information.

## Acknowledgements

The authors thank Yiqiao Wang (University of Oxford, UK) for contributions to degrader synthesis. AmpC and OXA-48 were kindly gifted by Professor Christopher Schofield (University of Oxford, UK). DegP and S210A-DegP expression plasmids were gifted by Professor Tim Clausen (Research Institute of Molecular Pathology, Austria) and Professor Seokhee Kim (Korea Advanced Institute of Science and Technology, South Korea). DSC was performed by David Staunton (Biophysics facility University of Oxford).

This research was supported by the Wellcome Trust (DPhil studentship 218514/Z/19/Z to E.K.T., and 317713/Z/24/Z to T.L.-H.), the Medical Research Council (MR/Y503484/1 to T.L.-H. and G.L.I.), European Research Council (ERC) under the European Union’s Horizon Europe research and innovation programme (grant agreement No. 101162143 awarded to G.L.I.), MR/W016672/1 (Medical Research Council career development award awarded to G.L.I), Royal Society of Chemistry (E22-1653035498 to T.L.-H. and G.L.I.), and the Medical and Life Sciences Translational Fund (CiC171/0014950 to T.L.-H. and G.L.I.) from the Medical Sciences Division, University of Oxford.

## Conflicts of Interests

The authors declare no competing interests.

## Data Availability Statement

The mass spectrometry proteomics data have been deposited to the ProteomeXchange Consortium via the PRIDE partner repository with the dataset identifier XXXXX

## Supplemental Figures

**Figure S1.**
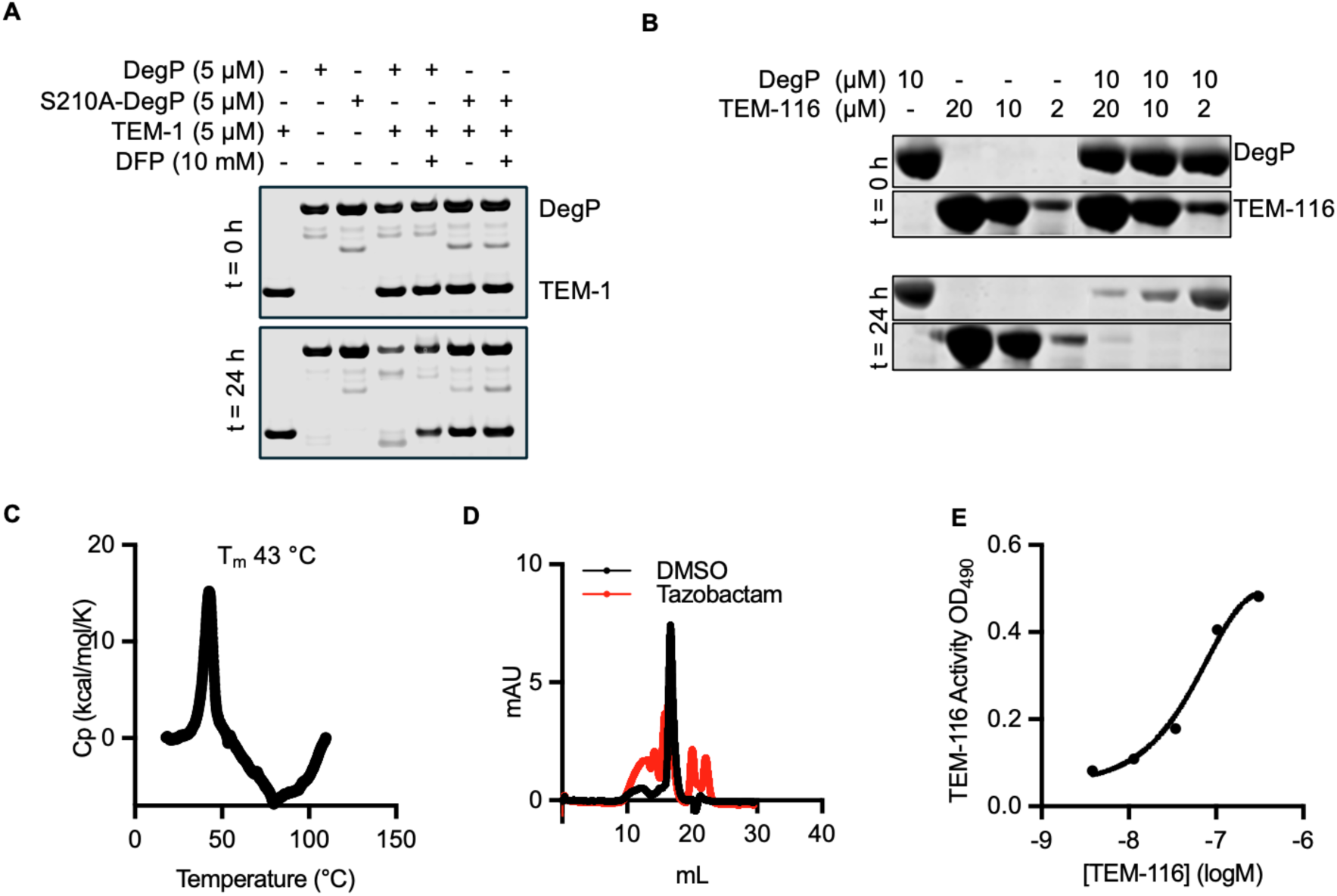
(A) DegP-mediated degradation of TEM-1 β-lactamase. (B) SDS-PAGE analysis showing increased TEM-116 concentration induced hyperactivation and autocleavage of DegP following 24 h incubation at 37 °C. (C) Differential scanning calorimetry analysis of TEM-116 showing melting temperature T_m_ = 43 °C, consistent with a folded protein. (D) Size-exclusion chromatogram of TEM-116 following incubation with DMSO or tazobactam for 4 h at 37 °C. Treatment with DMSO alone resulted in predominantly a single, monodisperse peak, representing folded protein, whereas tazobactam treatment induced multiple peaks. (E) Nitrocefin β-lactamase activity assay with increasing concentrations of TEM-116, demonstrating a functional protein. Data represent mean ± SEM (n = 3).

**Figure S2.**
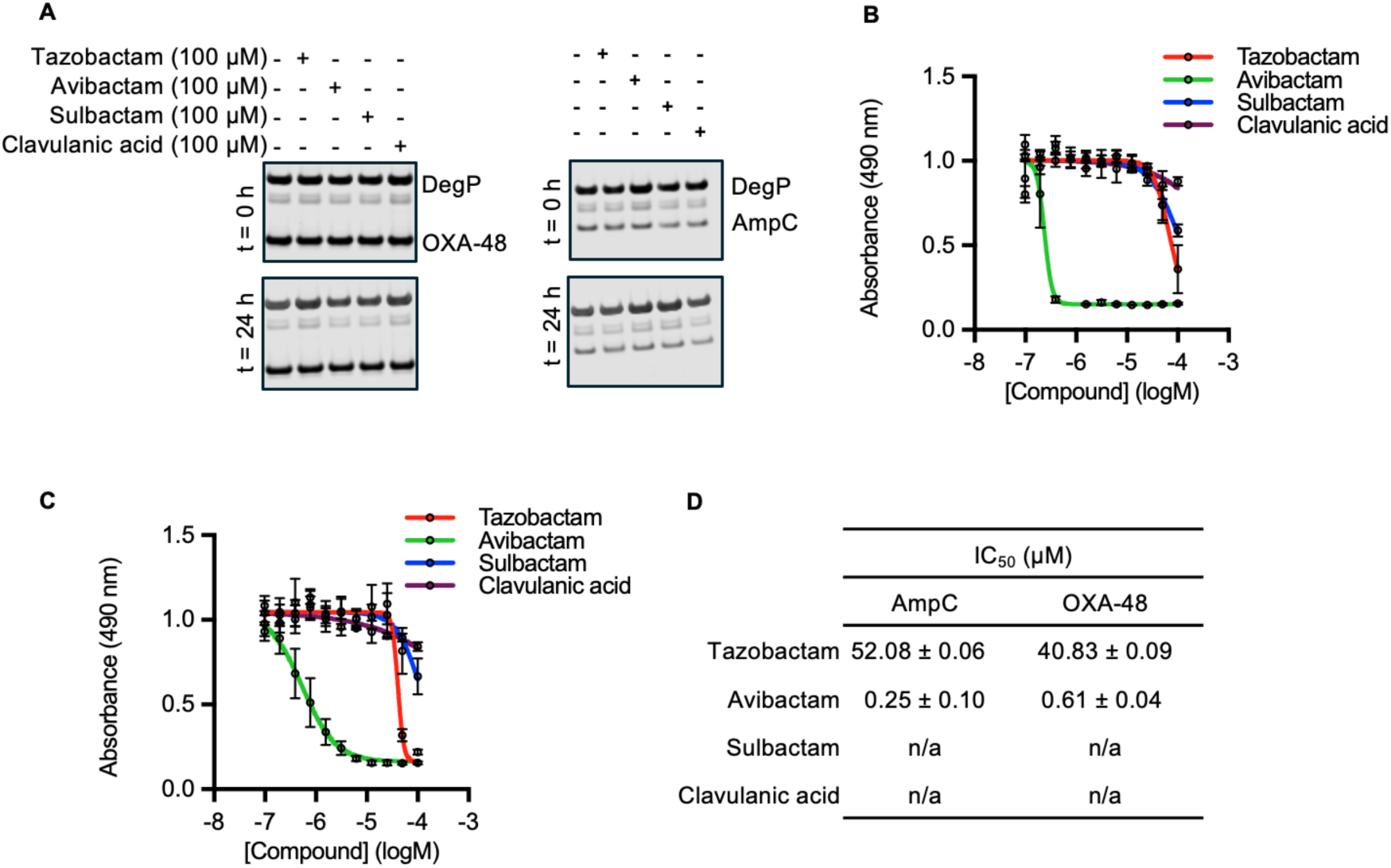
(A) SDS-PAGE analysis demonstrating a lack of effect of β-lactamase inhibitors on DegP-mediated degradation of OXA-48 and AmpC. (B) Dose-response analysis of β-lactamase inhibitors in AmpC nitrocefin assay. (C) Dose-response analysis of β-lactamase inhibitors in OXA-48 nitrocefin assay. (D) Respective IC_50_ values for inhibition of AmpC and OXA-48. Data represent mean ± SEM (n = 3).

**Figure S3.**
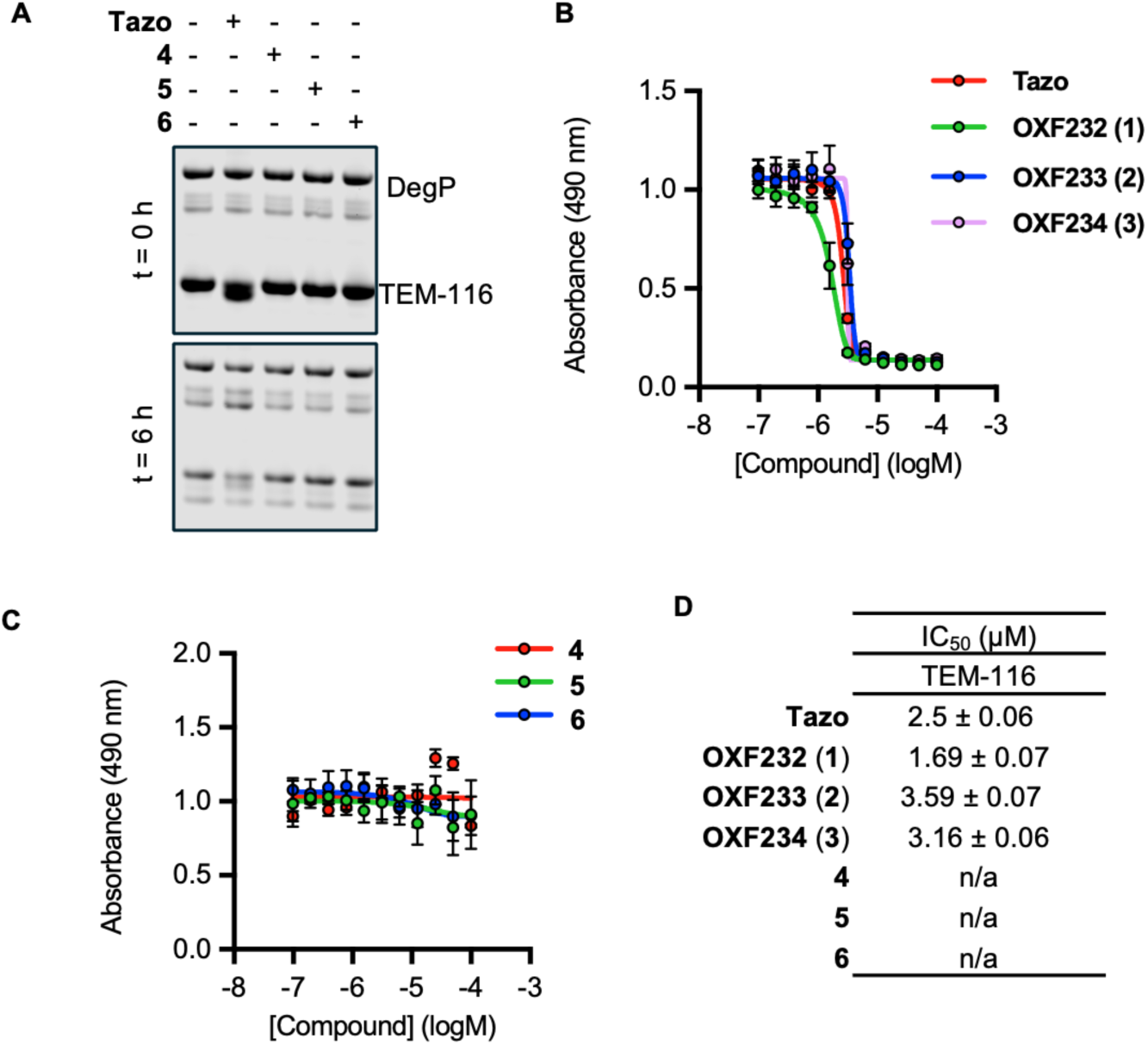
(A) SDS-PAGE analysis of the effect of amide-linked analogues **4**-**6** on DegP-mediated TEM-116 degradation. (B) Dose-response analysis of triazole-linked degraders **OXF232** (**1**), **OXF233** (**2**), and **OXF234** (**3**) in TEM-116 nitrocefin assay. (C) Dose-response analysis of amide-linked analogues **4**-**6** in TEM-116 nitrocefin assay (D) Respective IC_50_ values for inhibition of TEM-116. Data represent mean ± SEM (n = 3).

**Figure S4.**
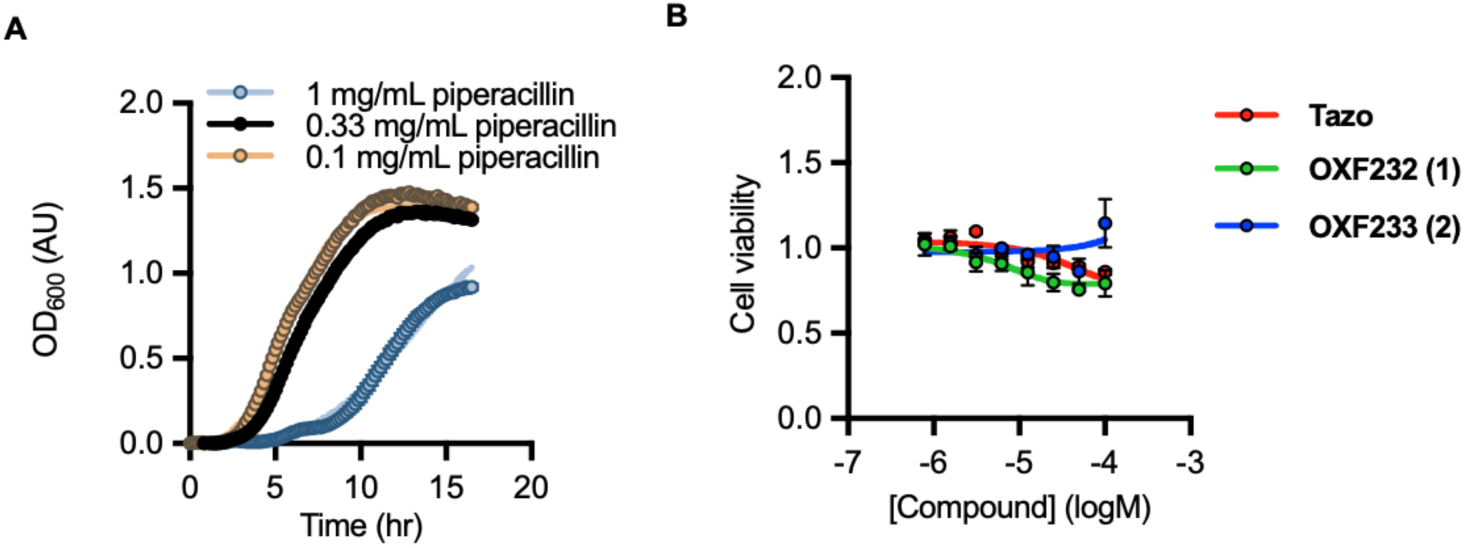
(A) Evaluation of piperacillin resistance in *E. coli* harbouring TEM-1 on the pK18-TEM1 plasmid. (B) Cytotoxicity testing of tazobactam (**Tazo**), **OXF232** (**1**), and **OXF233** (**2**) in HEK293T cells showing no major effects on cell growth. Data represent mean ± SEM (n = 3).

## Supplemental Schemes

**Scheme S1.**
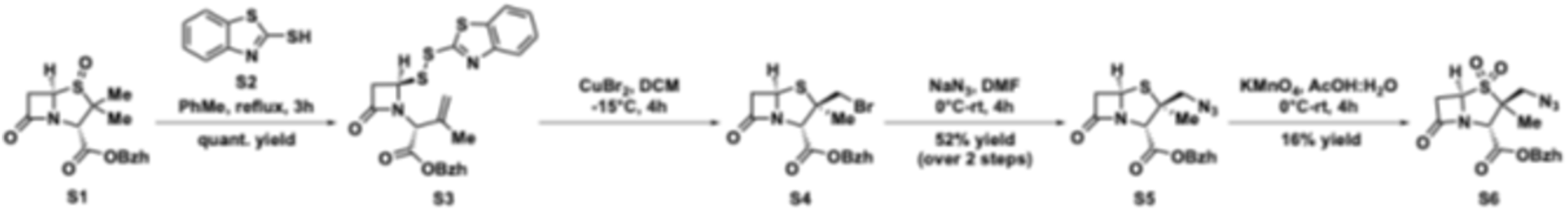
Synthesis of azide-containing tazobactam intermediate.[4]

## Supplemental Tables

**Table S1.**
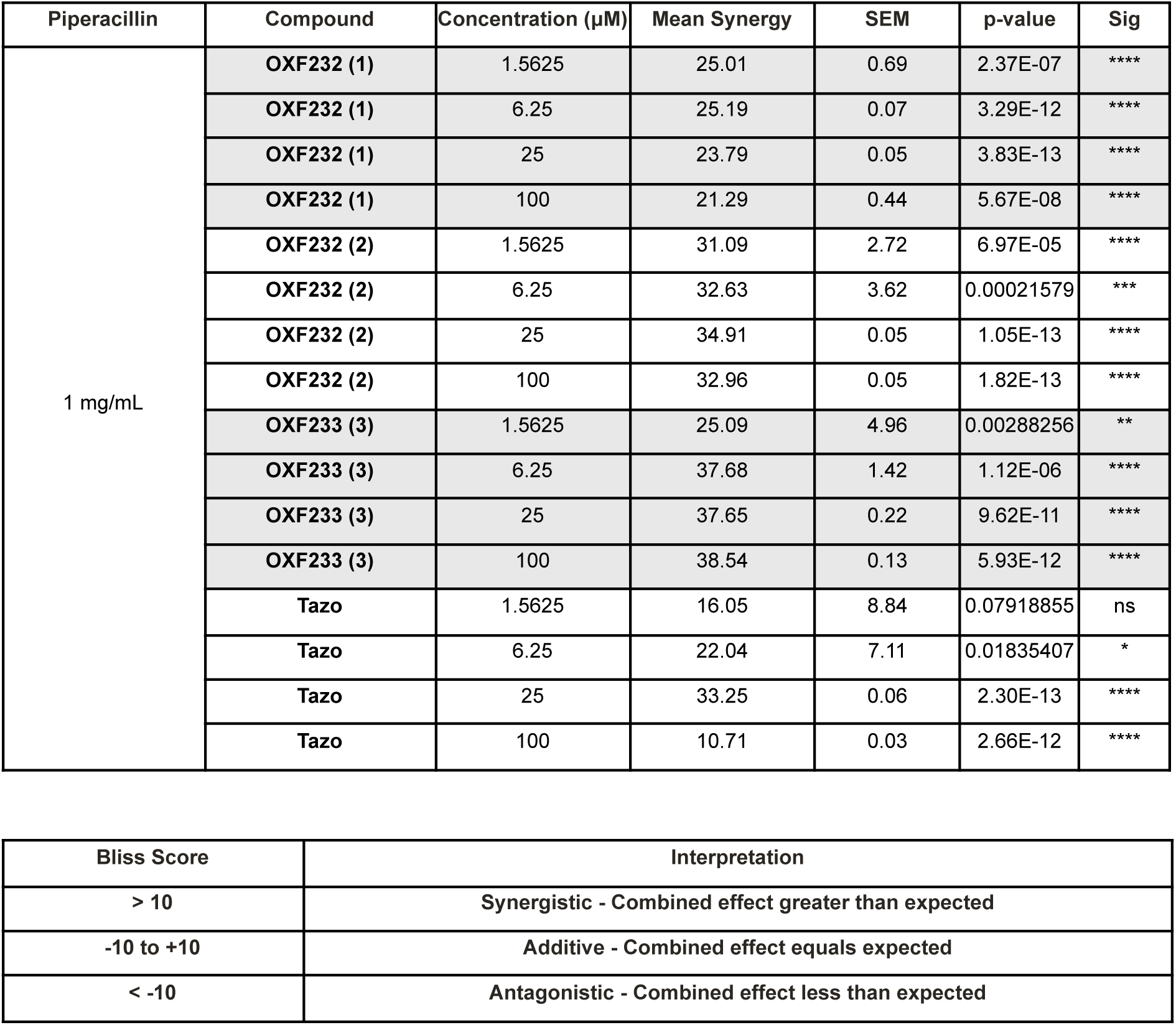
Bliss synergy score analysis of tazobactam (**Tazo**) and degraders **OXF232** (**1**), **OXF233** (**2**), and **OXF234** (**3**) for growth inhibition of *E. coli* harbouring TEM-1 on the pK18-TEM-1 plasmid, which contains the native constitutive promoter for TEM-1. Degraders **OXF232** (**1**), **OXF233** (**2**), and **OXF234** (**3**) showed improved synergy compared to tazobactam. Data represent mean ± SEM (n = 3).

**Table S2.** *In vitro* ADME properties of **OXF232 (1)** and **OXF233 (2)**.

| Assay | Reference / Benchmark | OXF232 (1) | OXF233 (2) |
| --- | --- | --- | --- |
| Solubility (kinetic, 0.1 M phosphate buffer, pH 7.4, $\mu\text{M}$) | >100 $\mu\text{M}$ = high solubility | >175 | >175 |
| LogD (octanol:buffer, 2:1) | Optimal range: 0–3 | -1.4 | -1.4 |
| **Human liver microsomes** |  |  |  |
| CLint ($\mu\text{L}\cdot\text{min}^{-1}\cdot\text{mg}^{-1}$) | Low < 9; High > 45 | <5.00 | <5.00 |
| Half-life (min) |  | >139 | >139 |
| **Mouse liver microsomes (CD-1)** |  |  |  |
| CLint ($\mu\text{L}\cdot\text{min}^{-1}\cdot\text{mg}^{-1}$) | Low < 10; High > 50 | <5.00 | <5.00 |
| Half-life (min) |  | >139 | >139 |

Note: Intrinsic clearance (CLint) values below the limit of quantification are reported as “<5.00 $\mu\text{L}\cdot\text{min}^{-1}\cdot\text{mg}^{-1}$ ”. Half-life values represent calculated lower bounds based on intrinsic clearance. Negative LogD values indicate high aqueous preference.

## Materials and Methods

### Biological methods

#### Protein Expression and Purification

**TEM-116.** The *tem-116* gene was cloned into a pET-28a(+) expression plasmid (GenScript) with an N-terminal His_10_-tag and 3C cleavage site and transformed into *E. coli* T7 Express (NEB). Overnight cultures were diluted 1:50 into fresh LB medium (Difco) supplemented with kanamycin (50 μg/mL) and grown at 37 °C with shaking until an OD_600_ of 0.7 was reached. Expression was induced with 1 mM IPTG and cultures were incubated overnight at 18 °C. Cells were harvested by centrifugation (5,000 × g, 30 min, 4 °C) and resuspended in lysis buffer (50 mM Tris-HCl pH 8.0, 300 mM NaCl). Cells were lysed by three passes through an Emulsiflex-C3 homogeniser (Avestin) at 15,000 psi and clarified by centrifugation (35,000 × g, 30 min, 4 °C). The supernatant was incubated with Ni-NTA resin (GE Healthcare #17531802) overnight at 4 °C. The resin was washed sequentially with wash buffer (50 mM Tris-HCl pH 8.0, 300 mM NaCl) containing 10, 20, 40, and 70 mM imidazole in 50 mM Tris-HCl pH 8.0, 300 mM NaCl. Bound protein was eluted with an elution buffer (50 mM Tris-HCl pH 8.0, 300 mM NaCl,250 mM imidazole). Eluted fractions were pooled and concentrated using a 10 kDa MWCO Amicon centrifugal filter (Millipore). For removal of the His-tag, TEM-116 was incubated with 3C protease (1:100, w/w) overnight at 4 °C, and passed over a Ni-NTA column to remove uncleaved protein, the cleaved His-tag, and His-tagged protease. Cleaved TEM-116 was further purified by size-exclusion chromatography on a Superdex 75 16/600 column (GE Healthcare) equilibrated in 50 mM Tris-HCl pH 8.0, 150 mM NaCl. Peak fractions were pooled, concentrated, flash-frozen in liquid nitrogen, and stored at −80 °C.

**DegP.** The *degP* gene was expressed from a pQE60 derivative with lacIQ and a C-terminal His_6_-tag (Spiess et al., 1999). The plasmid was transformed into *E. coli* T7 Express (NEB), and colonies were selected on LB-agar containing ampicillin (100 μg/mL). Overnight cultures were diluted 1:50 into LB with ampicillin, grown at 37 °C until an OD_600_ of 0.7 was reached. Expression was induced with 1 mM IPTG and cultures were incubated overnight at 18 °C. Cells were harvested (5,000 × g, 30 min, 4 °C), resuspended in lysis buffer (50 mM Tris-HCl pH 8.0, 300 mM NaCl, 10 mM imidazole), and lysed by Emulsiflex-C3 homogenization. The lysate was clarified (35,000 × g, 30 min) and incubated with Ni-NTA resin for 2 h at 4 °C. The resin was washed sequentially with wash buffer (50 mM Tris-HCl pH 8.0, 300 mM NaCl) containing 20 and 40 mM imidazole, and DegP was eluted with elution buffer (50 mM Tris-HCl pH 8.0, 300 mM NaCl, 250 mM imidazole). Eluted fractions were concentrated using a 30 kDa MWCO Amicon filter and further purified by size-exclusion chromatography on a Superdex 200 Increase 10/300 GL column (GE Healthcare) equilibrated in 50 mM Tris-HCl pH 8.0, 150 mM NaCl. Fractions corresponding to hexameric DegP were pooled, concentrated, flash-frozen in liquid nitrogen, and stored at −80 °C.

**S210A-DegP.** The catalytically inactive S210A-DegP mutant was expressed from the pCS21 plasmid, a pQE60 derivative carrying lacIQ and encoding DegPS210A with a C-terminal His_6_-tag (Spiess et al., 1999). The plasmid was transformed into an *E. coli* imp4213 Δ*degP* strain, and transformants were selected on LB-agar containing ampicillin (100 μg/mL). Overnight cultures were diluted 1:50 into LB medium with ampicillin, grown at 37 °C until an OD_600_ of 0.7 was reached. Expression was induced with 20 μM IPTG and cultures were incubated overnight at 18 °C. Cell harvesting, lysis and protein purification was performed as described for WT DegP, involving Ni-NTA affinity chromatography and Superdex 200 gel filtration. Fractions were pooled, concentrated, and stored at −80 °C.

#### *In vitro* degradation assay

Degradation assays were performed using purified β-lactamases (TEM-116, TEM-1, AmpC, and OXA-48) in the presence of DegP. Reactions were prepared in protease assay buffer (50 mM Tris-HCl, pH 8.0, 150 mM NaCl) in a total volume of 100 µL. Unless otherwise stated, all assays contained the β-lactamase (5 µM) and DegP (5 µM) and were incubated at 37 °C. Control reactions included (i) β-lactamase alone, (ii) β-lactamase with S210A-DegP (5 µM), and (iii) β-lactamase with DegP pre-incubated for 30 min on ice with diisopropyl fluorophosphate (DFP). For inhibitor and degrader testing, substrate proteins were pre-incubated for 10 min at 37 °C with the compounds prior to DegP addition. At the indicated timepoints, 10 µL aliquots were quenched with 4× SDS loading buffer, followed by heating at 95 °C for 5 min. Samples were analysed by SDS-PAGE (NuPAGE Bis-Tris 4–12%, Invitrogen) and visualised by Coomassie staining (LuBioScience).

#### Nitrocefin assay

β-lactamase (0.3 µM) were incubated with inhibitors at the specified concentrations for 1 h at 37°C in a total reaction volume of 50 uL PBS. Nitrocefin substrate was added (2.5 µL, 0.5 mg/mL), and absorbance at 490 nm was measured over two min. Data was analysed in Graphpad prism.

#### Sample preparation for mass spectrometry analysis

*E. coli* BW25113 was inoculated into LB (10 mL), and grown o/n at 37 °C with shaking (180 rpm). Cells were pelleted by centrifugation (4000 x g, 10 min, 4 °C), washed once with ice-cold PBS and resuspended in 1mL lysis buffer (50 mM Tris-HCl pH 8.0, 150 mM NaCl). Cells were lysed in a water-bath sonicator (ultra-high setting) for 5 min using 30 s on / 30 s off cycles, keeping samples on ice between. Lysates were clarified by centrifugation (≥20,000 × g, 15 min, 4 °C), and the protein concentration determined by BCA assay. Clarified lysates were combined with either (WT) DegP or S210A-DegP (final 5 µM) and incubated for 4 h at 37 °C. Following incubation, reactions were aliquoted to contain 150 µg input protein yielding n = 6 replicates per condition. Aliquots were flash-frozen in liquid nitrogen and stored at −80 °C until processing.

50 µg of protein lysate was subjected to SP3 bead digestion adapted from Hughes et al., 2014. Briefly, lysates were transferred to a 96-well plate (Nunc 29955). An equal volume of RIPA buffer (Pierce) supplemented with 10 mM TCEP (Thermo Scientific) and 20 mM chloracetamide (Sigma Aldrich) was added and samples incubated at 95 °C for 10 min. Two different SP3 beads (CYTIVA 4515210505025 and 65152105050250, 1 µL per sample) were washed three times in water and diluted 10-fold before adding to lysates. Acetonitrile was added to a final concentration of 70%, and the plate sealed and mixed on a plate shaker for 20 min. The plate was put on a 96-well magnet rack and the supernatant removed by inverting the plate, followed by two washes with 70% ethanol and a short wash with 100% acetonitrile. After complete evaporation of the acetonitrile, 100 µL of digestion buffer (Trypsin/LysC, 0.2 µg/sample, Thermo-Scientific, in freshly prepared 0.5 M urea, 50 mM Tris (pH 8.0), Sigma Aldrich) was added to each sample and incubated overnight at 37 °C. Digestion was stopped by addition of stop buffer (0.4% formic acid (FA), 4% acetonitrile) to a final concentration of 0.2% FA and 2% acetonitrile. The plate was put on a 96-well magnet rack and the peptide containing supernatant transferred to a new plate. Peptide concentration was determined using a colorimetric peptide assay kit (Pierce) and 250 µg of peptides were loaded on Evotip pure tips (Evosep) according to manufacturer’s instructions.

#### Mass spectrometry and data analysis

Evosep tips loaded with three samples of each treatment group were injected with an Evosep One LC system with an Evosep Endurance (15 cm x 150 µm, 1.9 µm) analytical column which was connected to a timsTOF Pro 2 system (Bruker). Samples were injected with 30 samples/day Evosep LC protocol. The HPLC mobile phase A was 0.1% FA, while mobile phase B contained 0.1% FA in acetonitrile. All mass spectra were collected in dia-PASEF mode.

Raw data were processed using directDIA in Spectronaut 20 (Biognosys). *E. coli* (strain B / BL21-DE3) reference protein database (UP000002032) with 4156 sequences was downloaded in October 2024 from UniProt. Volcano plot visualization was performed using Spectronaut standard settings. Unpaired t-test was applied with Qvalue set to confidence <0.05 and Log₂ ratio candidate filter set to 0.58.

#### C-terminal motif enrichment analysis

Proteins significantly decreased in the presence of WT DegP, compared to the untreated control, were identified from quantitative proteomic datasets. Proteins with a Log₂ fold change <-1 and adjusted p-value < 0.05 were classified as “degraded”, while non-significant hits were assigned to the “not degraded” group. Full-length sequences for each protein were retrieved from UniProt using accession numbers. The ten C-terminal amino acids were extracted using a custom R script. The following function was applied to each entry:

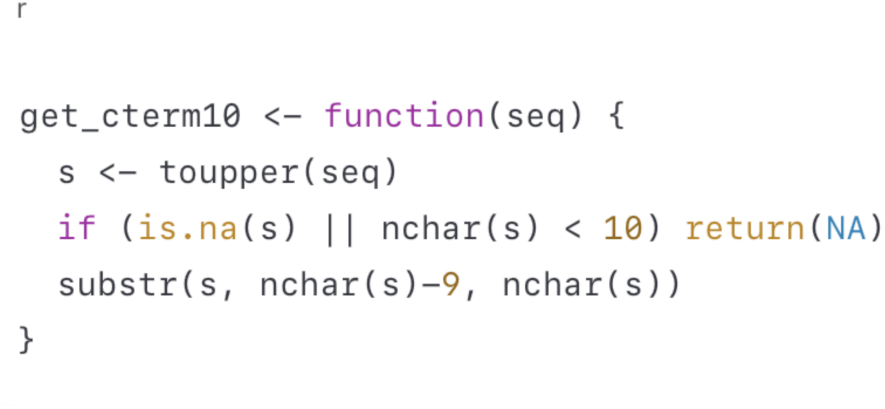

The resulting ten C-terminal residues were saved as plain text files for motif enrichment analysis. Amino acid frequency enrichment was visualized using TwoSampleLogo (v2.0; ^2^). The “degraded” dataset was uploaded as the positive set, and the “not degraded” group as the negative set. The “Protein” alphabet was selected with t-test as the statistical comparison method. Default parameters were used for all other settings. TwoSampleLogo plots were used to visualise residues enriched (above axis) or depleted (below axis) in the degraded group relative to controls. Statistical significance was assessed at p < 0.05 (uncorrected).

#### pK18-TEM-1 plasmid construction

The gene cassette containing *tem-1* with its native promoter was PCR-amplified from the *E. coli* clinical isolate E1N using the primers gaga<u>ggtacc</u>GAAATTGCTCATCAGCTCAG and atct<u>ggatcc</u>TTACCAATGCTTAATCAGTGAG (KpnI and BamHI restriction sites in underlined lowercase; genomic sequence in uppercase). To obtain pK18-TEM-1, the PCR amplicon was inserted into the multiple-cloning site of the pK18 vector via restriction digestion and ligation, and the resulting ligation mixture was transformed into *E. coli* NEB® 10-beta chemically competent cells. Transformants were selected at 37 °C on LB agar plates containing kanamycin (50 µg/mL) and the PCR-amplified DNA insert was verified by sequencing.

#### *E. coli* growth inhibition assay

The *E. coli* growth inhibition broth microdilution assay was performed according to CLSI methods M07-A11^4^ in 384-well plates (Greiner Bio-One, UK). Strains were grown overnight to stationary phase in LB and adjusted to OD_600_ of 0.005 prior to addition to the plate. Fourfold serial dilutions of test compounds were prepared in triplicate in LB to a final volume of 50 μL, with a no-compound growth control. Inoculum (25 μL) was added to each well to achieve a final OD_600_ of 0.0025, excluding a no-inoculum sterility control, which had only LB added (final volume 50 μL, 1% (*v/v*) DMSO). Cells were incubated at 37 °C in a BMG LABTECH FLUOstar Omega 18 h, with OD_600_ readings taken every 10 min. OD_600_ measurements were background-corrected against a no-inoculum control and normalised to a DMSO control for dose-response analysis.

#### Differential Scanning Calorimetry (DSC)

Differential Scanning Calorimetry (DSC) experiments were performed on a VP PEAQ DSC instrument (Malvern Panalytical, UK). Samples and buffer blanks were measured using a thermal ramp of 200 °C/h between 10 °C and 110 °C at a pressure of 55 psi. The protein concentration studied on the instrument was 20 μM. Heat capacity versus temperature data were analysed by Microcal PEAQ-DSC software v1.53 and the T_m_ and enthalpy of unfolding calculated.

#### Pharmacokinetic assays

Kinetic solubility, equilibrium solubility, human liver microsome stability and mouse liver microsome stability analysis was performed by Concept Life Science.

#### Cell toxicity assay

HEK293T viability was quantified using the CellTiter-Glo Luminescent Cell Viability Assay (Promega, Cat. no. G7570) according to the manufacturer’s instructions. Briefly, HEK293T cells were seeded in white, flat-bottom 96-well plates at a density of 10,000 cells per well in 100 µL Complete growth medium overnight at 37 °C, 5% CO₂. Cells were treated with compounds or vehicle control (0.1% v/v DMSO) for 24 h at 37 °C. Following treatment, CellTiter-Glo reagent (100 µL) was added directly to each well, and plates were mixed on an orbital shaker for 2 min to induce cell lysis, followed by a 10 min incubation at room temperature. Luminescence was measured using a microplate luminometer. Background signal from wells containing medium and CellTiter-Glo reagent only was subtracted from all readings. Viability in each well was expressed as a percentage of the mean luminescence of vehicle-treated control wells on the same plate.

### Strains and Plasmids

The following strains and plasmids were used in this study:

| Bacterial strain | Description | Resistance Markers (Concentration) | Source |
| --- | --- | --- | --- |
| *E. coli* BW25113 K12<sup>[1]</sup> | Wild-type *E. coli* K-12 derivative | n/a | Baba, T *et al.*, 1999 |
| T7 express *E. coli* | Chemically competent BL21 derived *E. coli* | n/a | NEB (USA) |
| pCS20 | pQE60 derivative with *lacI*<sup>Q</sup>, *degP* (C-terminal His<sub>6</sub>-tag) | Ampicillin (100 µg/mL) | Spiess *et al.*, 1999 |
| pCS21 | pQE60 derivative with *lacI*<sup>Q</sup>, *degPS210A* (C-terminal His<sub>6</sub>-tag) | Ampicillin (100 µg/mL) | Spiess *et al.*, 1999 |
| *E. coli* imp4213 ΔdegP | Hyperpermeable *imp4213* background with *degP* deletion | n/a | Coli Genetic Stock Center (Yale University) |
| *E. coli* imp4213 (RFM795) ΔdegP | Hyperpermeable *imp4213* background with *degP* deletion | n/a | Coli Genetic Stock Center (Yale University) |
| *E. coli* MG1655 ΔampC | *E. coli* MG1655 ΔampC expressing TEM1A from a high-copy number plasmid | Kanamycin (50 μg/mL) | This study |

### Chemical methods

#### General Information

Commercially available chemicals were purchased from Sigma-Aldrich, Fluorochem or Novabiochem line (Merck KGaA) and used as received. All reactions were performed in an argon atmosphere unless otherwise stated. Solvents used for chromatography were HPLC grade and used as received. **Analytical thin-layer chromatography (TLC)** was performed on silica gel 60 F_254_ pre-coated plates (Merck, UK) and visualized under UV light (254 nm). **Automated solid-phase peptide synthesis** was performed using Liberty Blue 2.0 automated microwave peptide synthesizer (CEM Corporation). **Purifications by preparative high-performance liquid chromatography (HPLC)** were performed on a SP20AP system (Shimadzu) equipped with a Prominence SPD-20A UV/Vis detector and FRC-10A fraction collector. Separations were performed using an XBridge BEH C18 OBDä preparative column (130 Å, 5mm, 19 x 150mm), eluting with a gradient of 30-95% acetonitrile in water + 0.1% FA at a flow rate of 17 mL/min over 16 min. **Nuclear Magnetic Resonance (NMR)** spectra were recorded on a AVIII HD Nanobay 400 MHz NMR spectrometer (Bruker), equipped with a 5 mm z-gradient multinuclear BBFO probe using the software TopSpin (version 3, Bruker BioSpin) or on a Avance NEO 600 MHz NMR spectrometer (Bruker), equipped with a 5 mm BB-F/^1^H helium-cooled cryoprobe using the software TopSpin (version 4, Bruker BioSpin). **Chemical shifts (δ)** for ^1^H and ^13^C NMR spectra are given in ppm relative to residual solvent chemical shifts converted to the TMS scale (CDCl_3_: δH = 7.26 ppm, δC = 77.16 ppm, DMSO-*d*_6_: δH = 2.50 ppm, δC = 39.52 ppm). Data are reported as follows: chemical shift (δ), multiplicity, coupling constants (*J*) in Hertz, and integrated intensity. **High-Resolution Mass Spectrometry (HRMS)** analyses were recorded on an ACQUITY I-Class PLUS UPLC System (Waters, USA) coupled to an ACQUITY RDa mass spectrometer (Waters, USA) equipped with an ESI probe, in positive ion mode. **Ultra-performance liquid chromatography (UPLC) coupled to HRMS** analyses were performed on an ACQUITY UPLC liquid chromatograph (Waters, UK) system coupled direct to a HRMS Waters Xevo G2-XS QTOF (Waters, UK) equipped with StepWave^TM^ ion-transfer optics and an ESI LockSprayä source. The stationary phase was equipped with ACQUITY Premier BEH C18 column (130 Å, 1.7 μm, 2.1 × 50 mm). The mobile phase was acetonitrile and water with FA as additive with a gradient of 5-95% acetonitrile in water (+0.1% FA) over 10 min. UPLC-HRMS analysis confirmed all final compounds possessed purities ≥95%.

#### Synthesis of azide-containing tazobactam intermediate:[4]

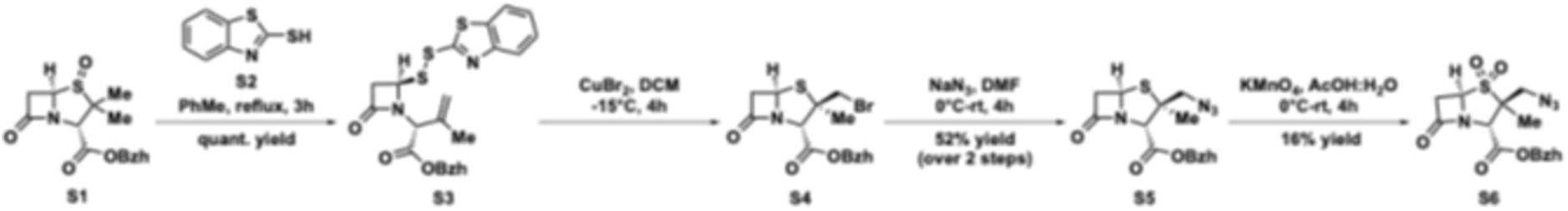

#### Synthesis of the azetidinone disulfide intermediate (S3)

2-Mercaptobenzothiazole **S2** (881 mg, 1.01 equiv., 5.27 mmol) was added to a stirred solution of benzhydryl (2S,5R)-3,3-dimethyl-7-oxo-4-thia-1-azabicyclo[3.2.0]heptane-2-carboxylate 4-oxide **S1** (20.0 mL, 0.27 M). The resulting reaction mixture was refluxed for 3 h. After that time, the reaction was concentrated *in vacuo*, MeOH (20 mL) was added to remove residual PhMe and the solid was washed 3x with minimal amount of DCM to afford the crude benzhydryl (R)-2-((R)-2-(benzo[d]thiazol-2-yldisulfaneyl)-4-oxoazetidin-1-yl)-3-methylbut-3-enoate **S3** (2.91 g, 5.46 mmol, quantitative yield) as a yellow gum, which was used in the next step without further purification. ^1^H NMR (400 MHz, CDCl_3_) δ 7.90 – 7.85 (m, 1H), 7.80 – 7.75 (m, 1H), 7.47 – 7.43 (m, 1H), 7.38 – 7.27 (m, 11H), 6.90 (s, 1H), 5.39 (dd, *J* = 5.0, 2.3 Hz, 1H), 5.17 – 5.12 (m, 1H), 5.03 – 4.98 (m, 1H), 4.92 – 4.89 (m, 1H), 3.44 (dd, *J* = 15.4, 5.0 Hz, 1H), 3.21 (dd, *J* = 15.4, 2.3 Hz, 1H), 1.91 (s, 3H). HRMS (ESI+) calculated for [C_29_H_24_N_2_O_3_S_3_]^+^ [M+H]^+^: 533.1047, found: 533.1022. Rf (20% EtOAc in PE) = 0.31.

#### Synthesis of the bromomethylpenicillin intermediate (S4)

Copper (II) bromide (1.40 g, 1.15 equiv., 6.28 mmol) was added to a solution of benzhydryl (R)-2-((R)-2-(benzo[d]thiazol-2-yldisulfaneyl)-4-oxoazetidin-1-yl)-3-methylbut-3-enoate **S3** (2.91 g, 1 equiv., 5.46 mmol) in anhydrous DCM (26.00 mL) at −10 °C. The resulting reaction mixture was stirred at 0 °C for 4 h. The reaction mixture was then diluted with DCM (100 mL). Celite was added, stirred for 10 min, and filtered off. The filtrate was washed with 1% NaHCO_3_ solution (50 mL), followed by water (50 mL), and brine (50 mL). The organic layer was dried over Na_2_SO_4_ and concentrated *in vacuo* to afford a crude residue containing the product benzhydryl (2S,3R,5R)-3-(bromomethyl)-3-methyl-7-oxo-4-thia-1-azabicyclo[3.2.0]heptane-2-carboxylate **S4** (1.48 g, 3.32 mmol, 61% yield) as a yellow gum, which was used immediately in the next step. ^1^H NMR (400 MHz, CDCl_3_) δ 7.39 – 7.29 (m, 10H), 6.94 (s, 1H), 5.44 (dd, *J* = 4.1, 1.8 Hz, 1H), 5.20 (s, 1H), 3.62 (dd, *J* = 16.1, 4.1 Hz, 1H), 3.56 (s, 2H), 3.14 (dd, *J* = 16.1, 1.8 Hz, 1H), 1.36 (s, 3H).

#### Synthesis of the azidomethylpenicillin intermediate (S5)

The residue from previous step was dissolved in anhydrous DMF (40 mL) and cooled to 0 °C. Then, a solution of sodium azide (533 mg, 1.5 equiv., 8.19 mmol) in water (12 mL) was added dropwise and the resulting reaction was gradually warmed to room temperature and stirred for 4 h. After that time, the reaction was diluted with water (40 mL) and extracted with EtOAc (100 mL). The organic layer was sequentially washed with water (150 mL), and brine (8 x 50 mL), dried over Na_2_SO_4_ and concentrated *in vacuo*to afford the product benzhydryl (2S,3S,5R)-3-(azidomethyl)-3-methyl-7-oxo-4-thia-1-azabicyclo[3.2.0]heptane-2-carboxylate **S5** (1.17 g, 2.86 mmol, 52% yield) as a yellow gum. ^1^H NMR (400 MHz, CDCl_3_) δ 7.42 – 7.31 (m, *J* = 4.6 Hz, 10H), 6.94 (s, 1H), 5.38 (dd, *J* = 4.1, 1.8 Hz, 1H), 4.84 (s, 1H), 3.60 (dd, *J* = 16.0, 4.1 Hz), 3.47 (s, 2H), 3.12 (dd, *J* = 16.0, 1.8 Hz, 1H), 1.24 (s, 3H). HRMS analysis (ESI+) calculated for [C_21_H_20_N_4_O_3_SNa]^+^ [M+Na]^+^: 431.1148, found: 431.1168.

#### Synthesis of the azidomethylpenicillin-dioxide intermediate (S6)

Under ice-water bath conditions, KMnO_4_ (905 mg, 2 equiv., 5.73 mmol) was added portion wise to a solution of benzhydryl (2S,3S,5R)-3-(azidomethyl)-3-methyl-7-oxo-4-thia-1-azabicyclo[3.2.0]heptane-2-carboxylate **S5** (1.17 g, 1 equiv., 2.86 mmol) in AcOH:H_2_O (19:1, 23 mL). The reaction was gradually warmed to room temperature and stirred for 4 h. After that time, the mixture was cooled with an ice-batch and hydrogen peroxide solution (30 wt. % in H_2_O) was added dropwise until the solution turned white. The mixture was diluted with ice-water (100 mL) and extracted with DCM (3x 50 mL). The organic layer was washed with 1% NaHCO_3_ solution (50 mL), brine (50 mL), dried over anhydrous Na_2_SO_4_, and concentrated *in vacuo* to afford a crude residue. The residue was purified by preparative HPLC and lyophilised to afford the benzhydryl (2S,3S,5R)-3-(azidomethyl)-3-methyl-7-oxo-4-thia-1-azabicyclo[3.2.0]heptane-2-carboxylate 4,4-dioxide **S6** (330 mg, 0.749 mmol, 16 % yield) as a white powder. ^1^H NMR (400 MHz, CDCl_3_) δ 7.41 – 7.31 (m, 10H), 6.97 (s, 1H), 4.63 (s, 1H), 4.60 (dd, *J* = 4.1, 2.2 Hz, 1H), 3.91 (d, *J* = 13.3 Hz, 1H), 3.75 (d, *J* = 13.3 Hz, 1H), 3.53 (dd, *J* = 16.3, 4.2 Hz, 1H), 3.47 (dd, *J* = 16.2, 2.2 Hz, 1H), 1.18 (s, 3H). HRMS analysis (ESI+) calculated for [C_21_H_20_N_4_O_5_SNa]^+^ [M+Na]^+^: 463.1047, found: 463.1074.

#### Synthesis of degraders 1-3 linked via the triazole ring

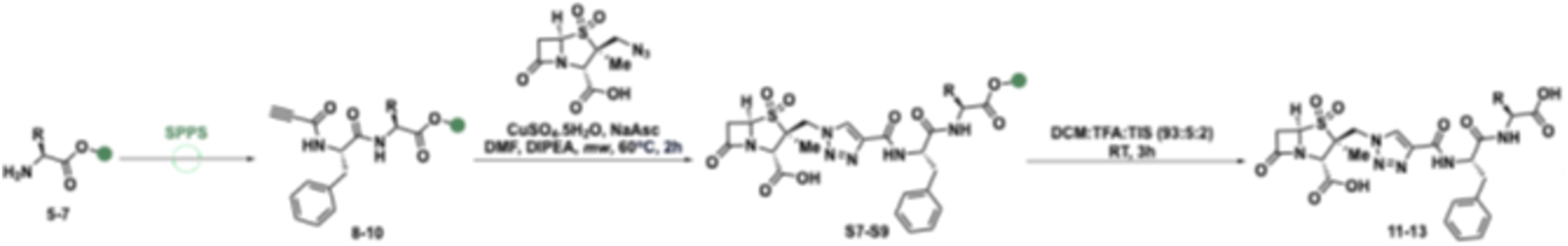

#### General method for one-pot solid-phase peptide synthesis and click reaction

The solid-phase peptide synthesis started with a respective preloaded 2-ClTrt resin (0.1 mmol scale) and sequential coupling of the amino acid Fmoc-Phe-OH and propiolic acid were performed using a 5-fold excess of each reagent. Coupling reactions were performed at 50 °C for 10 min under microwave irradiation using 0.5 M DIC solution in DMF (5-fold excess) and 1.0 M OxymaPure solution with 0.1 M DIPEA in DMF (5-fold excess). Fmoc-deprotection was performed using 20% piperidine (v/v) in DMF (4 mL per cycle) at 75 °C for 3 min under microwave irradiation. Initial and final Fmoc deprotection steps were not required. After completion of the automated sequence of amide couplings and Fmoc-deprotection, the peptide-2-ClTrt resin was functionalised via click reaction. Benzhydryl (2S,3S,5R)-3-(azidomethyl)-3-methyl-7-oxo-4-thia-1-azabicyclo[3.2.0]heptane-2-carboxylate 4,4-dioxide **5** (0.2 M solution in DMF, 0.1 mmol), CuSO_4_.5H_2_O (32 mg, 0.13 mmol), sodium ascorbate (50 mg, 0.25 mmol), DIPEA (35 mL, 0.1 mmol) and DMF (3 mL) were added to the reaction vessel containing the resin-bound peptide and the resulting reaction mixture was heated at 60 °C for 2 h under microwave irradiation. The final resin-bound peptide was obtained as a DMF suspension. The suspension was transferred to a Poly-prep® chromatography column, the solvent was removed, the resin washed with DCM (15 mL) and diethyl ether (15 mL) and dried under an argon flow. The resin-bound peptide was treated with a cold solution of DCM:TFA:TIS (93:5:2; 5 mL). The resulting suspension was mixed at room temperature for 3 h. The solution was collected and concentrated under an argon flow until the volume decreased to 2 mL. The resulting residue was precipitated by addition of diethyl ether (8 mL) and isolated by centrifugation. The crude residue was purified by preparative HPLC and lyophilised to afford the respective product.

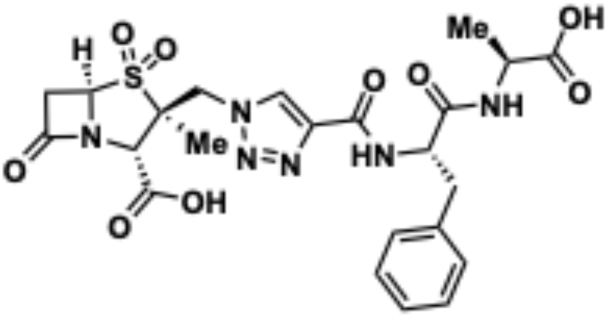

**OXF232** (**1**) was obtained in 12% yield (6.73 mg, 12.0 μmol) as a white solid from H-Ala-2-ClTrt resin (0.137g, 0.73 mmol/g loading capacity).

**^1^H NMR (400 MHz, DMSO-*d_6_*) δ** 8.61 (d, *J* = 7.3 Hz, 1H), 8.55 (s, 1H), 8.41 (d, *J* = 8.6 Hz, 1H), 7.42 –7.20 (m, 5H), 5.32 (d, *J* = 15.3 Hz, 1H), 5.25 (dd, *J* = 4.4, 1.6 Hz, 1H), 5.04 (d, *J* = 15.2 Hz, 1H), 4.85 (td, *J* = 9.1, 4.2 Hz, 1H), 4.78 (s, 1H), 4.34 (p, *J* = 7.3 Hz, 1H), 3.77 (dd, *J* = 16.4, 4.5 Hz, 1H), 3.38 (dd, *J* = 16.4, 1.6 Hz, 1H), 3.19 (dd, *J* = 13.8, 4.2 Hz, 1H), 3.10 (dd, *J* = 13.9, 9.5 Hz, 1H), 1.45 (s, 3H), 1.40 (d, *J* = 7.3 Hz, 3H).

**^13^C NMR (151 MHz, DMSO-*d_6_*) δ** 174.0, 171.4, 170.6, 167.7, 159.0, 142.0, 137.7, 129.3, 128.4, 128.1, 126.3, 64.3, 61.9, 60.2, 53.5, 50.5, 47.6, 40.1, 37.7, 37.5, 17.2, 15.8.

**UPLC–HRMS (ESI+)** RT = 3.22 min (100% purity); m/z [M + H]⁺ calculated for C_23_H_27_N_6_O_9_S^+^ 563.1555, found 563.1555.

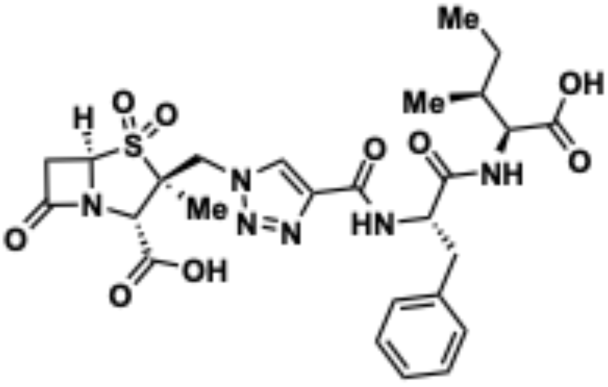

**OXF234** (**2**) was obtained in 11% yield (6.79 mg, 11.2 μmol) as a white solid from H-Ile-2-ClTrt resin (0.141g, 0.71 mmol/g loading capacity).

**^1^H NMR (400 MHz, DMSO-*d_6_*) δ** 8.42 – 8.35 (m, 2H), 8.27 (d, *J* = 8.5 Hz, 1H), 7.32 – 7.11 (m, 5H), 5.14 (s, 1H), 5.04 – 5.00 (m, 1H), 4.88 – 4.78 (m, 1H), 4.23 (dd, *J* = 8.5, 5.8 Hz, 2H), 4.18 (s, 1H), 3.62 – 3.53 (m, 2H), 3.19 (d, *J* = 16.2 Hz, 1H), 3.12 – 3.06 (m, 1H), 3.06 – 2.98 (m, 1H), 1.81 (s, 1H), 1.48 – 1.39 (m, 1H), 1.30 (s, 3H), 0.91 – 0.82 (m, 6H).

**^13^C NMR (151 MHz, DMSO-*d_6_*) δ** 172.8, 171.5, 170.9, 158.9, 142.0, 137.6, 129.3, 128.2, 128.0, 126.3, 64.8, 61.8, 56.3, 53.6, 50.8, 40.1, 37.4, 37.1, 36.5, 36.5, 24.7, 15.7, 15.5, 11.3.

**UPLC–HRMS (ESI+)** RT = 3.73 min (96% purity); m/z [M + H]⁺ calculated for C_26_H_33_N_6_O_9_S^+^ 605.2024, found 605.2068.

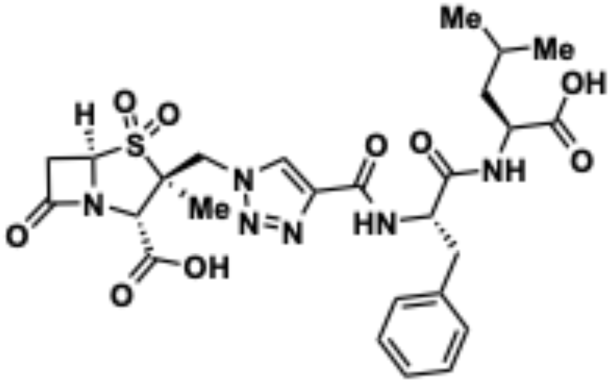

**OXF234** (**3**) was obtained in 5% yield (2.90 mg, 4.80 μmol) as a white solid from H-Leu-2-ClTrt resin (0.154g, 0.65 mmol/g loading capacity).

**^1^H NMR (400 MHz, DMSO-*d_6_*) δ** 8.43 – 8.38 (m, 2H), 8.32 (d, *J* = 8.6 Hz, 1H), 7.32 – 7.12 (m, 5H), 5.13 (d, *J* = 7.3 Hz, 2H), 5.08 – 5.03 (m, 1H), 4.80 – 4.74 (m, 1H), 4.27 (s, 1H), 3.64 – 3.56 (m, 1H), 3.21 (d, *J* = 16.3 Hz, 1H), 3.10 (dd, *J* = 13.7, 4.3 Hz, 1H), 3.02 (dd, *J* = 13.8, 9.4 Hz, 1H), 1.69 – 1.63 (m, 1H), 1.60 – 1.52 (m, 2H), 1.32 (s, 3H), 0.91 (d, *J* = 6.5 Hz, 3H), 0.85 (d, *J* = 6.5 Hz, 3H).

**^13^C NMR (151 MHz, DMSO-*d_6_*) δ** 173.9, 171.5, 170.8, 158.9, 142.0, 137.6, 129.3, 128.2, 128.0, 126.3, 64.7, 61.8, 53.5, 50.7, 50.3, 40.1, 37.4, 37.2, 24.3, 22.9, 21.4, 15.7.

**UPLC–HRMS (ESI+)** RT = 3.80 min (100% purity); m/z [M + H]⁺ calculated for C_26_H_33_N_6_O_9_S^+^ 605.2024, found 605.2068.

#### Synthesis of compounds 4-6 linked via the carboxylic acid group

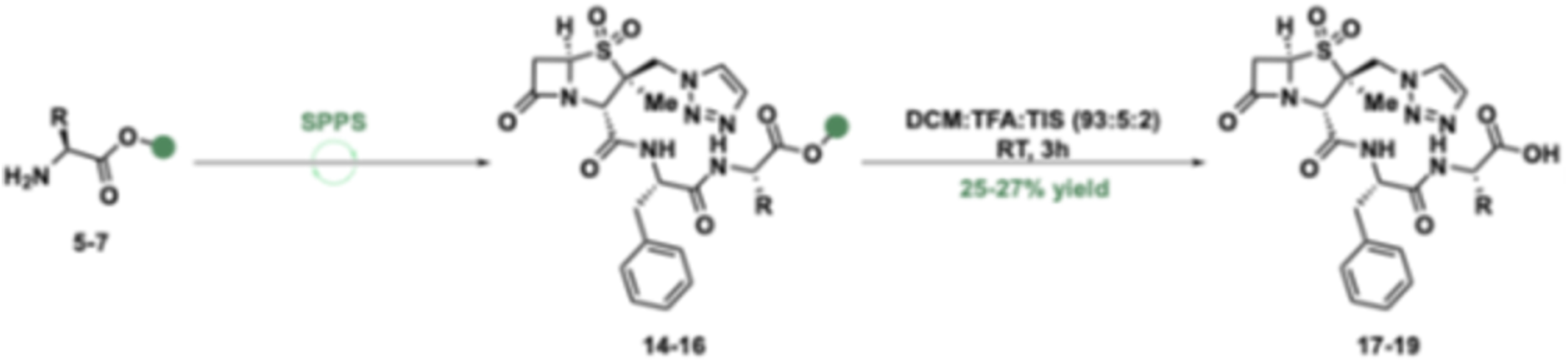

#### General method for SPPS

The solid-phase peptide synthesis started with a respective preloaded 2-ClTrt resin (0.1 mmol scale) and sequential coupling of the amino acid Fmoc-Phe-OH and tazobactam were performed using a 5-fold excess of each reagent. Coupling reactions were performed at 90 °C for 2 min under microwave irradiation using 0.5 M DIC solution in DMF (5-fold excess) and 1.0 M OxymaPure solution with 0.1 M DIPEA in DMF (5-fold excess). Fmoc-deprotection was performed using 20% piperidine (v/v) in DMF (4 mL per cycle) at 90 °C for 1 min under microwave irradiation. Initial and final Fmoc-deprotection steps were not required. After completion of the automated sequence of amide couplings and Fmoc-deprotection, the final resin-bound peptide was obtained as a DMF suspension. The suspension was transferred to a Poly-prep® chromatography column, the solvent was removed, the resin washed with DCM (15 mL) and diethyl ether (15 mL) and dried under an argon flow. The resin-bound peptide was treated with a cold solution of DCM:TFA:TIS (93:5:2; 5 mL). The resulting suspension was mixed at room temperature for 3 h. The solution was collected and concentrated under an argon flow until the volume decreased to 2 mL. The resulting residue was precipitated by the addition of diethyl ether (8 mL) and isolated by centrifugation. The crude residue was purified by preparative HPLC and lyophilised to afford the respective product.

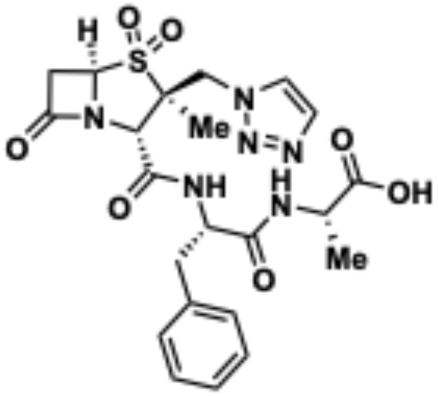

Compound **4** was obtained in 27% yield (14.13 mg, 27.25 μmol) as a white solid from H-Ala-2-ClTrt resin (0.127g, 0.79 mmol/g loading capacity).

**UPLC–HRMS (ESI+)** RT = 3.53 min (100% purity); m/z [M + H]⁺ ^+^ 520.1729, found 520.1949.

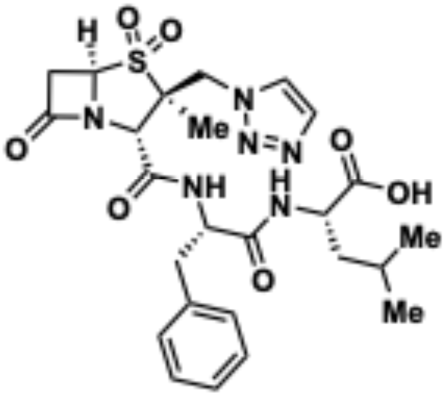

Compound **5** was obtained in 25% yield (13.73 mg, 24.49 μmol) as a white solid from H-Leu-2-ClTrt resin (0.154g, 0.65 mmol/g loading capacity).

**UPLC–HRMS (ESI+)** RT = 4.10 min (100% purity); m/z [M + H]⁺ ^+^ 561.2126, found 561.2084.

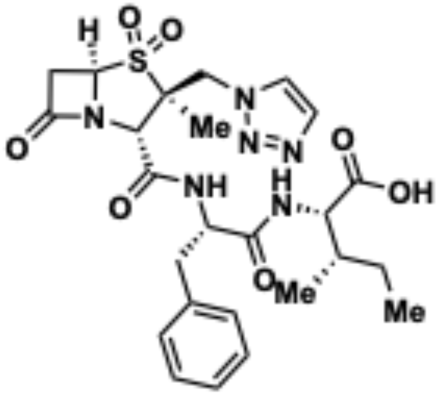

Compound **6** was obtained in 25% yield (13.98 mg, 24.94 μmol) as a white solid from H-Ile-2-ClTrt resin (0.141g, 0.71 mmol/g loading capacity).

**UPLC–HRMS (ESI+)** RT = 4.03 min (100% purity); m/z [M + H]⁺ ^+^ 561.2126, found 561.2084.

#### Synthesis of dipeptide *N*-Ac-Phe-Ala-OH

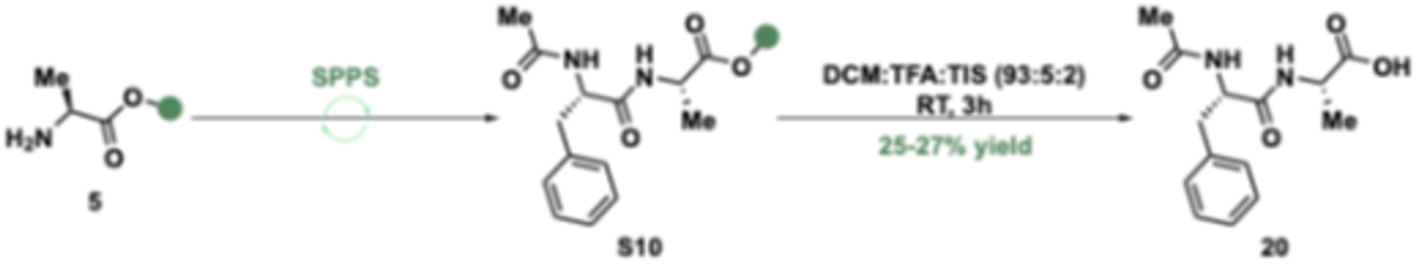

The solid-phase peptide synthesis started with preloaded H-Ala-2-ClTrt resin (0.1 mmol, 0.137g, 0.73 mmol/g loading capacity) and the coupling of *N*-Ac-Phe-OH was performed using a 5-fold excess of each reagent. The coupling reaction was performed at 50 °C for 10 min under microwave irradiation using 0.5 M DIC solution in DMF (5-fold excess) and 1.0 M OxymaPure solution with 0.1 M DIPEA in DMF (5-fold excess). After the automated amide coupling, the final peptide-resin bounded was obtained as a DMF suspension. The suspension was transferred to a Poly-prep® chromatography column, the solvent was removed, the resin washed with DCM (15 mL) and diethyl ether (15 mL) and dried under an argon flow. The resin-bound peptide was treated with a cold solution of DCM:TFA:TIS (93:5:2; 5 mL). The resulting suspension was mixed at room temperature for 3 h. The solution was collected and concentrated under an argon flow. The crude residue was purified by preparative HPLC and lyophilised to afford the expected dipeptide acetyl-L-phenylalanyl-L-alanine (**7**) in 48% yield (13.34 mg, 47.93 μmol) as a white solid. **^1^H NMR (400MHz, DMSO-*d_6_*) δ** 8.31 (d, J=7.3Hz, 1H), 8.06 (d,J=8.6Hz,1H), 7.29–7.14 (m, 5H), 4.54–4.48 (m, 1H), 4.18 (p, J=7.3Hz, 1H), 3.00 (dd, J=13.9, 3.9 Hz, 1H), 2.69 (dd, J=13.9, 10.3 Hz, 1H), 1.73 (s, 3H), 1.28 (d, J = 7.3 Hz, 3H). **^13^C NMR (101 MHz, DMSO-*d_6_*) δ** 174.1, 171.4, 169.3, 138.1, 129.2, 128.1, 126.3, 53.7, 47.7, 40.4, 37.7, 22.5, 17.2.

**UPLC–HRMS (ESI+)** RT = 2.92 min (100% purity); m/z [M + H]⁺ calculated for C_14_H_19_N_2_O_4_^+^ 279.1339, found 279.1362.

#### NMR and UPLC-HRMS analysis of final compounds

(2*S*,3*S*,5*R*)-3-((4-(((*S*)-1-(((*S*)-1-Carboxyethyl)amino)-1-oxo-3-phenylpropan-2-yl)carbamoyl)-1*H*-1,2,3-triazol-1-yl)methyl)-3-methyl-7-oxo-4-thia-1-azabicyclo[3.2.0]heptane-2-carboxylic acid 4,4-dioxide (OXF232, 1)

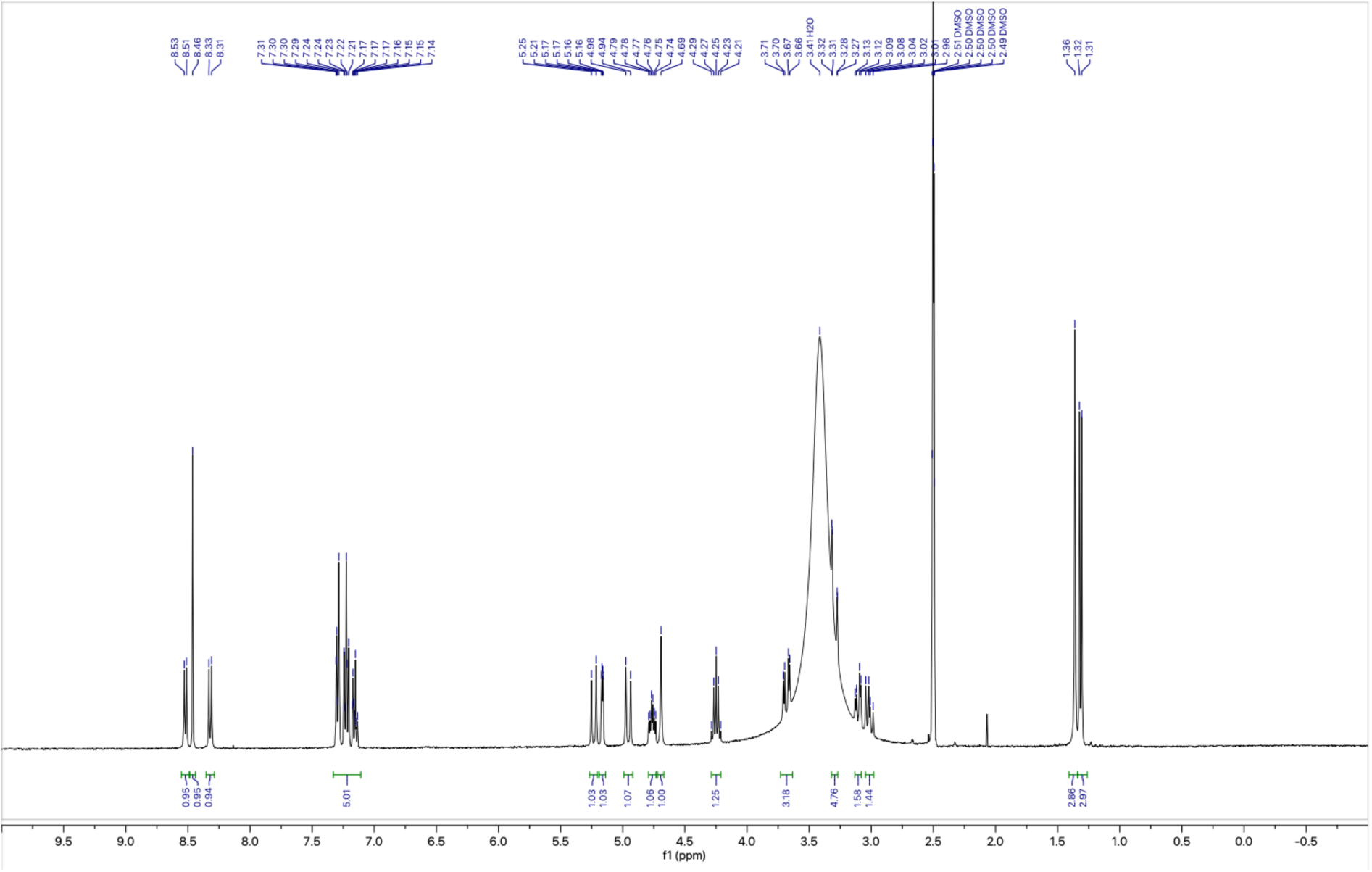

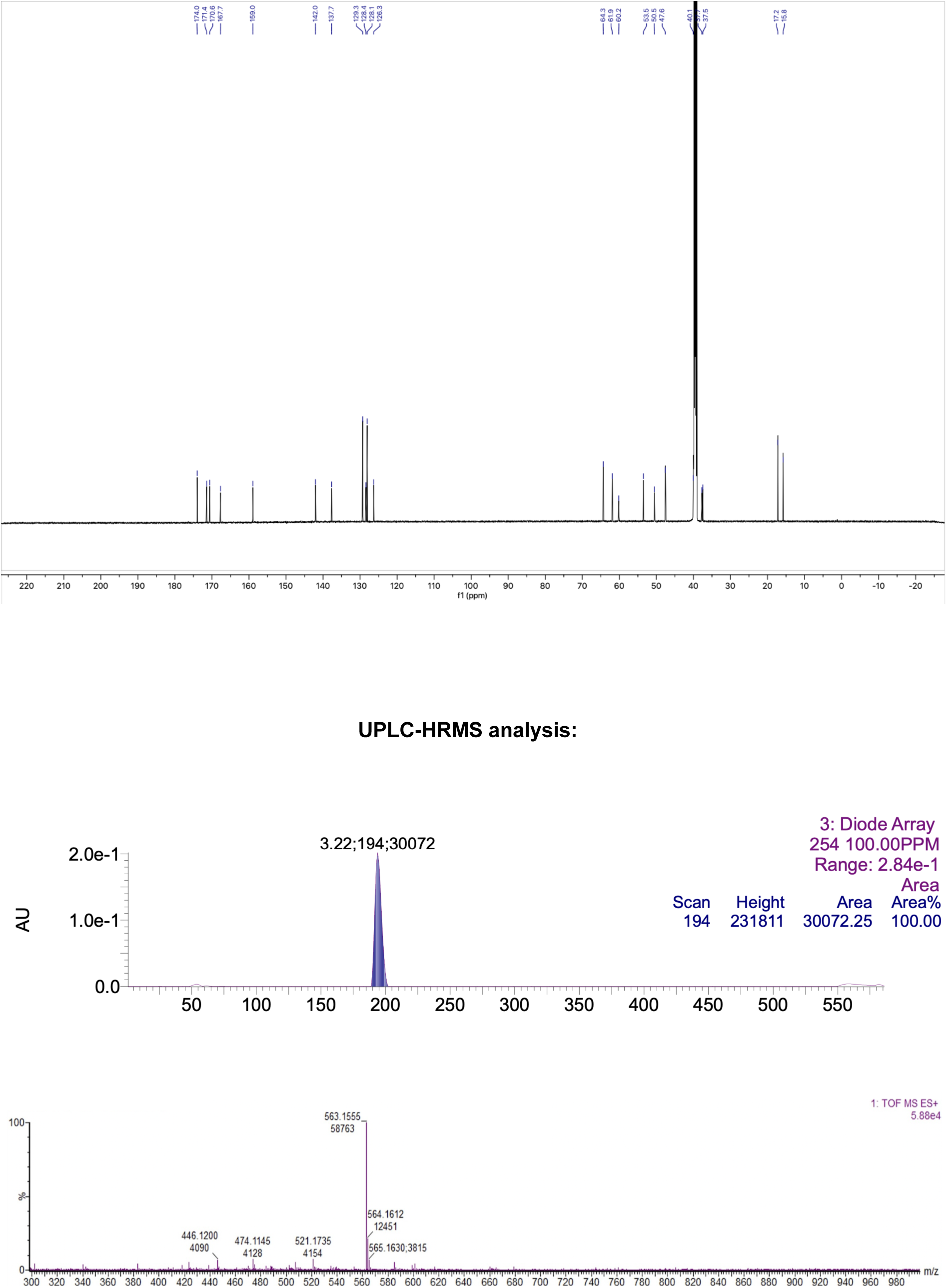

(2*S*,3*S*,5*R*)-3-((4-(((*S*)-1-(((1*S*,2*S*)-1-Carboxy-2-methylbutyl)amino)-1-oxo-3-phenylpropan-2-yl)carbamoyl)-1*H*-1,2,3-triazol-1-yl)methyl)-3-methyl-7-oxo-4-thia-1-azabicyclo[3.2.0]heptane-2-carboxylic acid 4,4-dioxide (OXF233, 2)

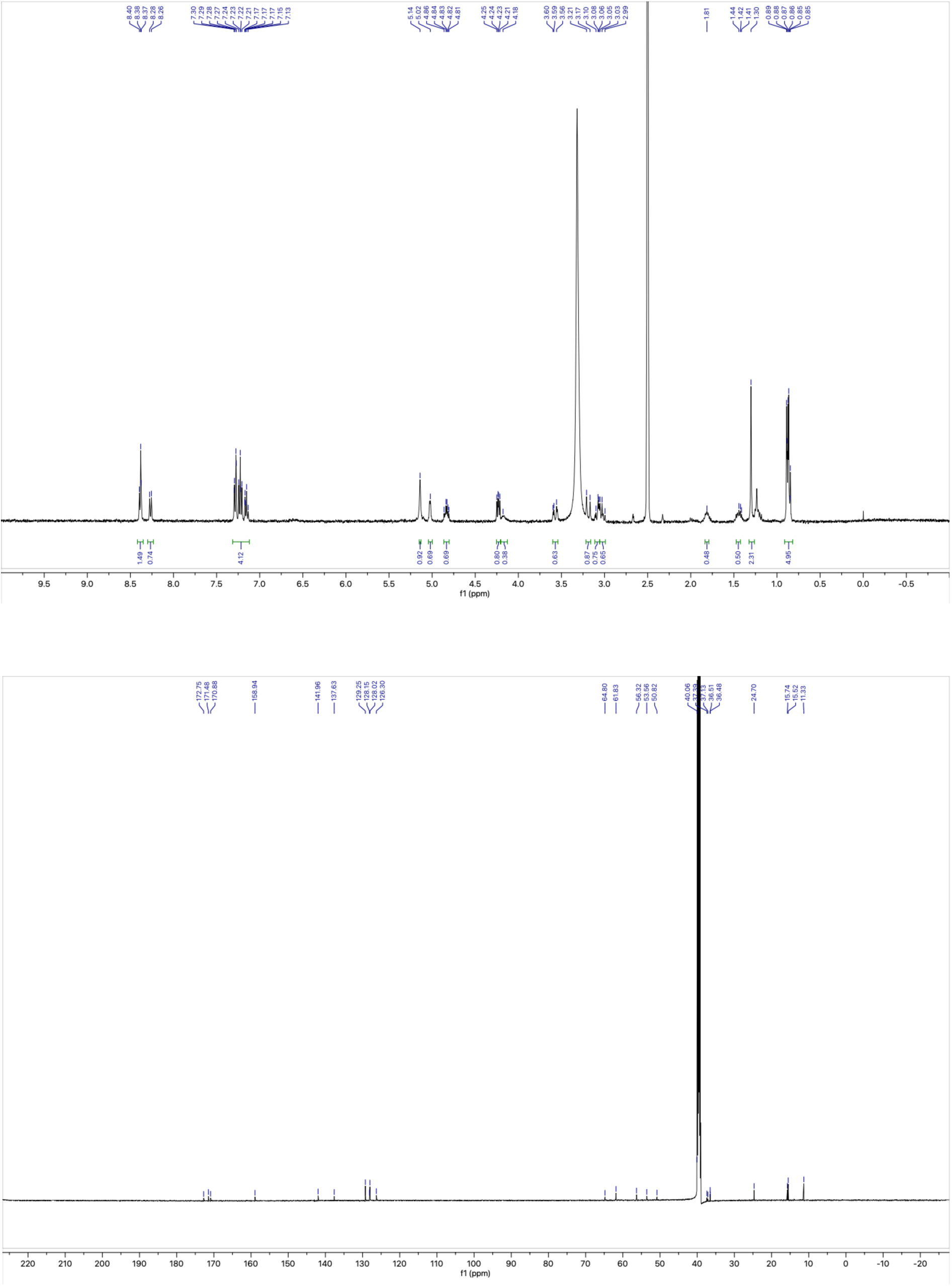

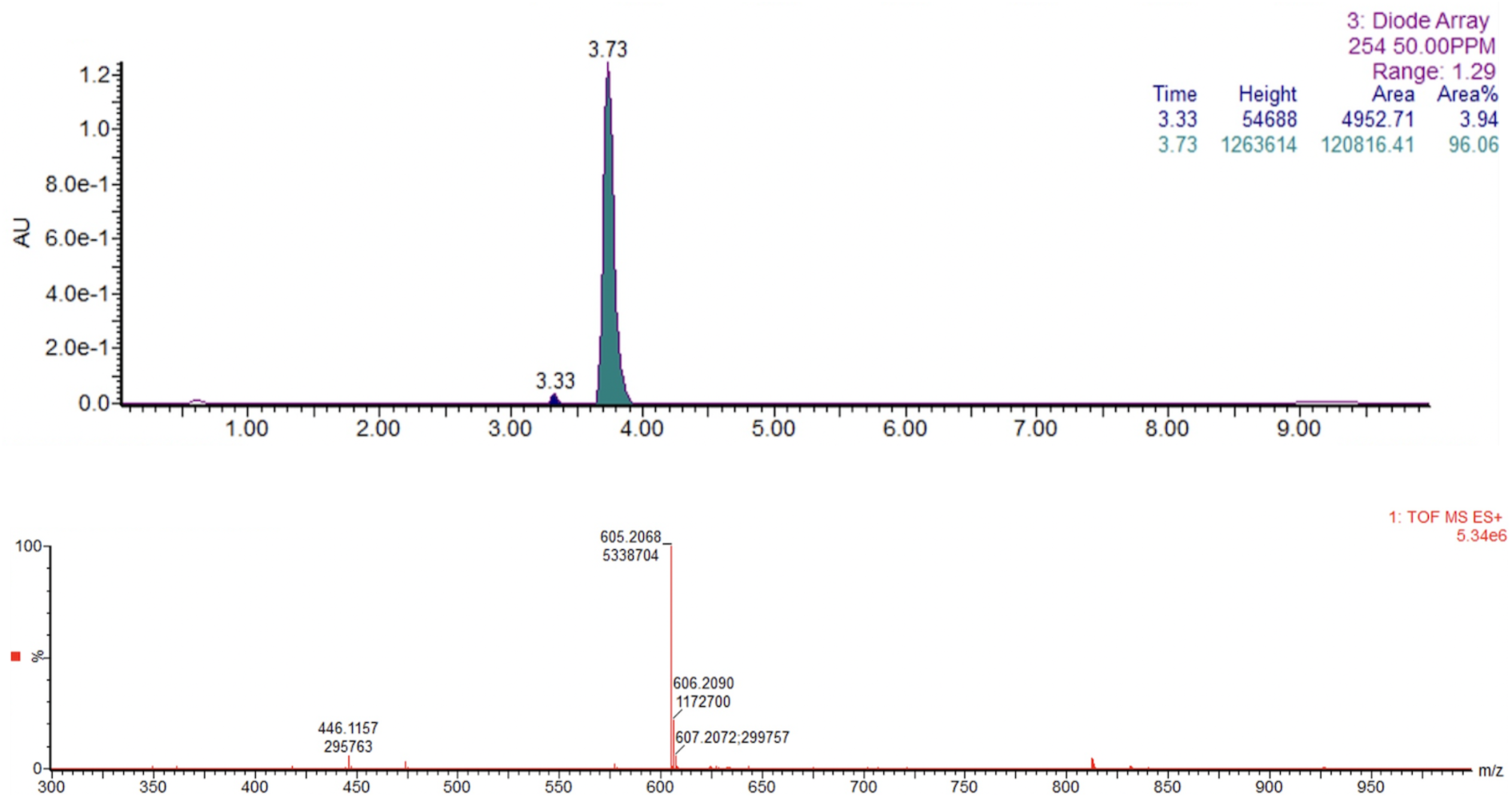

(2*S*,3*S*,5*R*)-3-((4-(((*S*)-1-(((*S*)-1-Carboxy-3-methylbutyl)amino)-1-oxo-3-phenylpropan-2-yl)carbamoyl)-1*H*-1,2,3-triazol-1-yl)methyl)-3-methyl-7-oxo-4-thia-1-azabicyclo[3.2.0]heptane-2-carboxylic acid 4,4-dioxide (OXF234, 3)

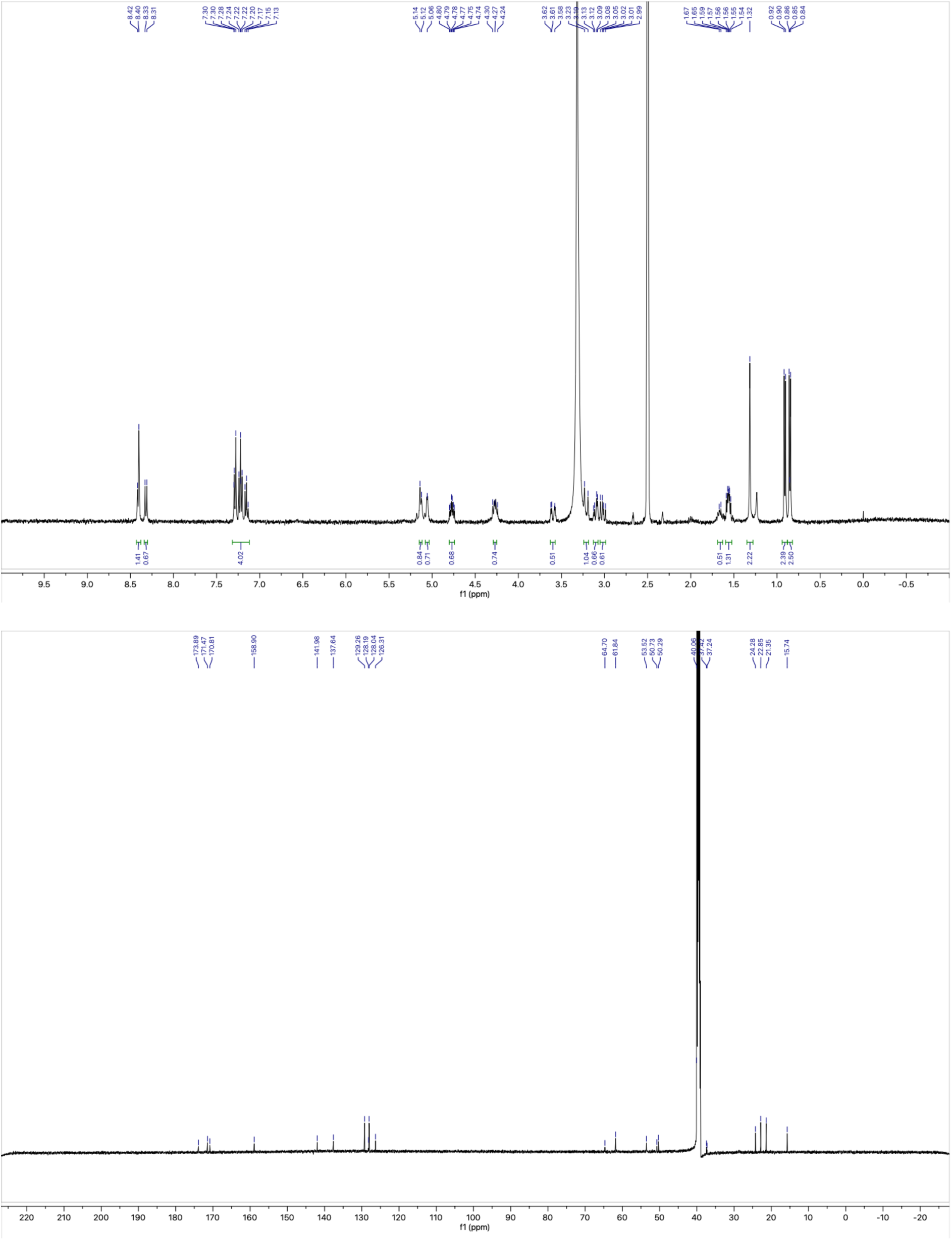

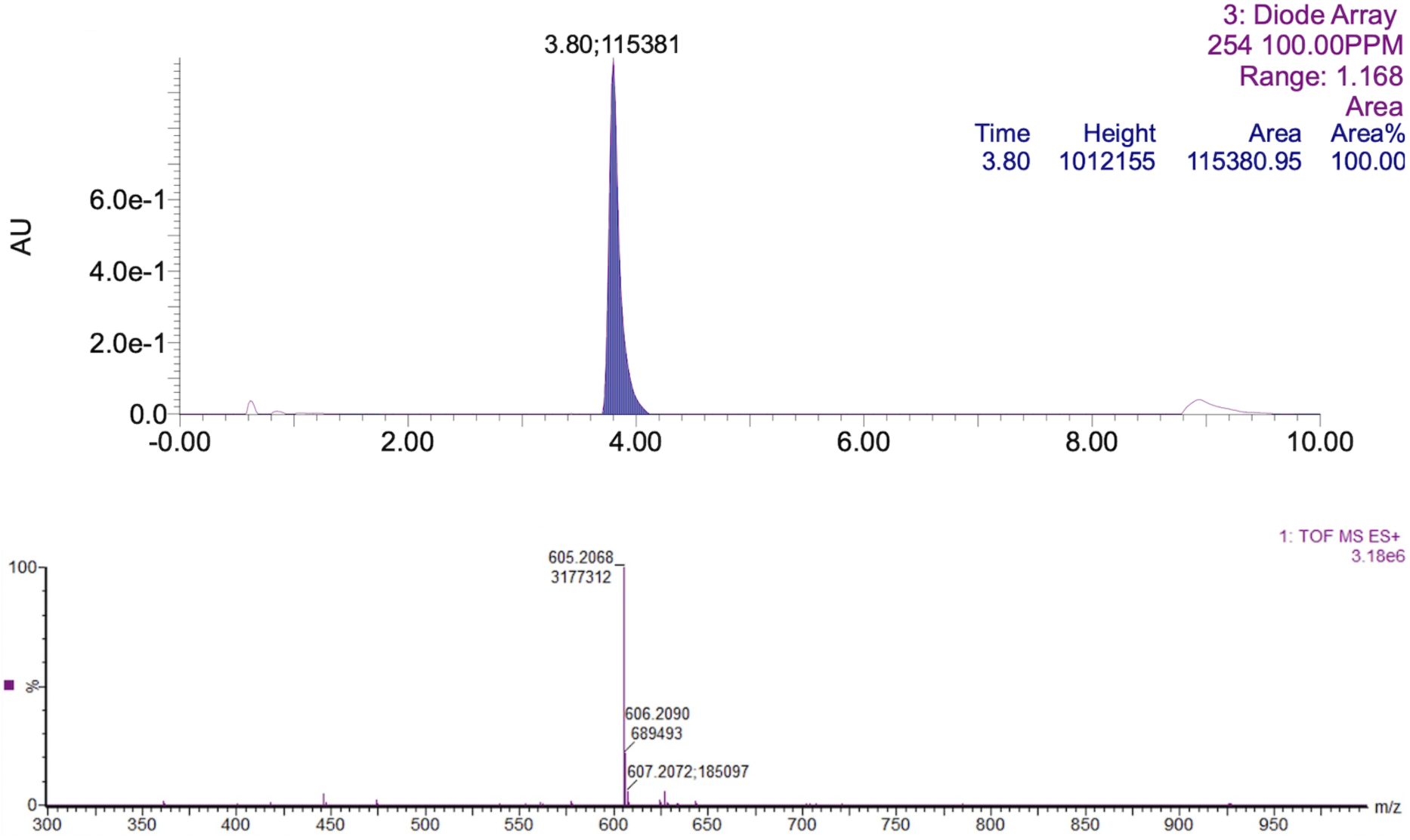

((2S,3S,5R)-3-((1H-1,2,3-Triazol-1-yl)methyl)-3-methyl-4,4-dioxido-7-oxo-4-thia-1-azabicyclo[3.2. **0]heptane-2-carbonyl)-L-phenylalanyl-L-alanine (4)**

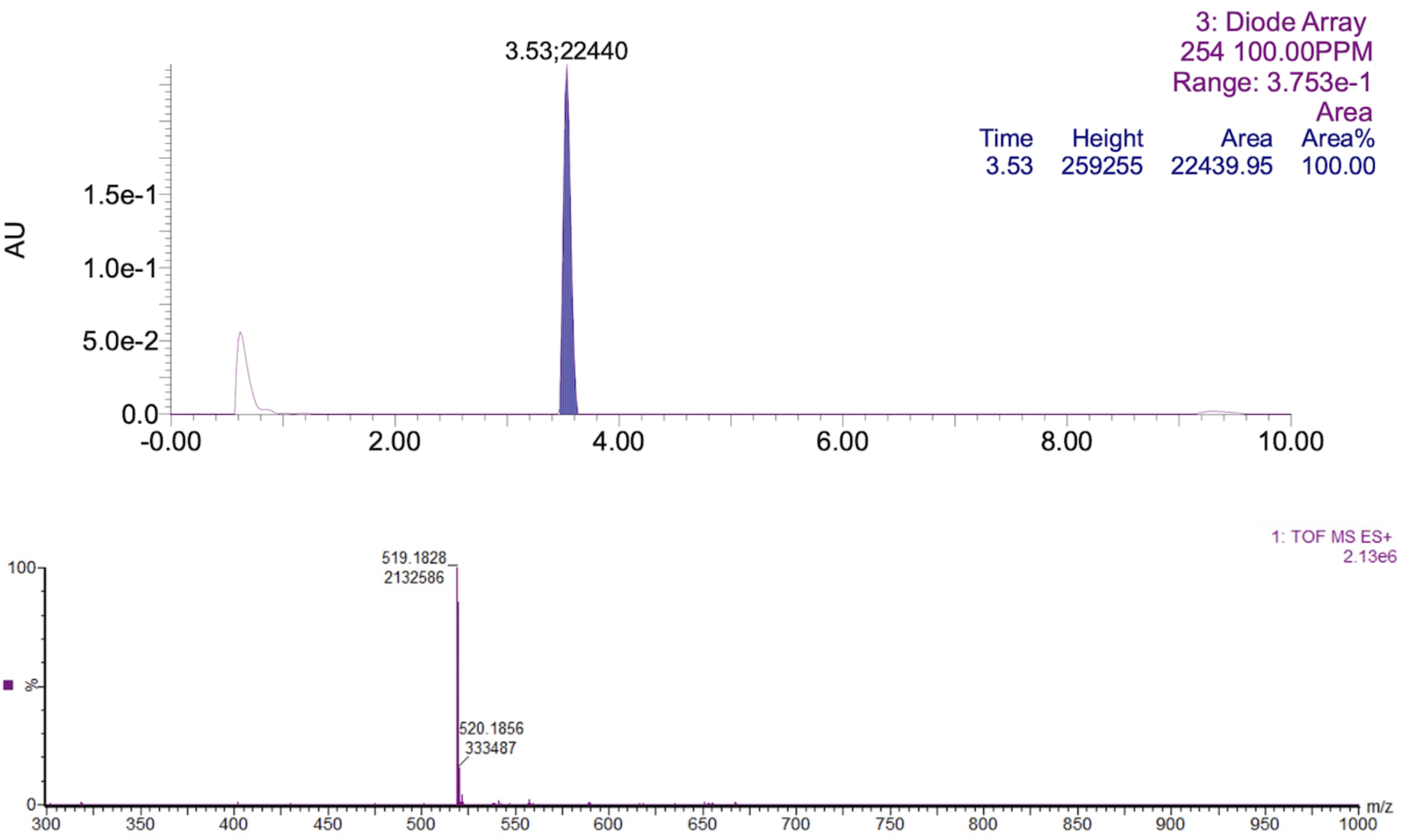

((2S,3S,5R)-3-((1H-1,2,3-Triazol-1-yl)methyl)-3-methyl-4,4-dioxido-7-oxo-4-thia-1-azabicyclo[3.2. **0]heptane-2-carbonyl)-L-phenylalanyl-L-isoleucine (5)**

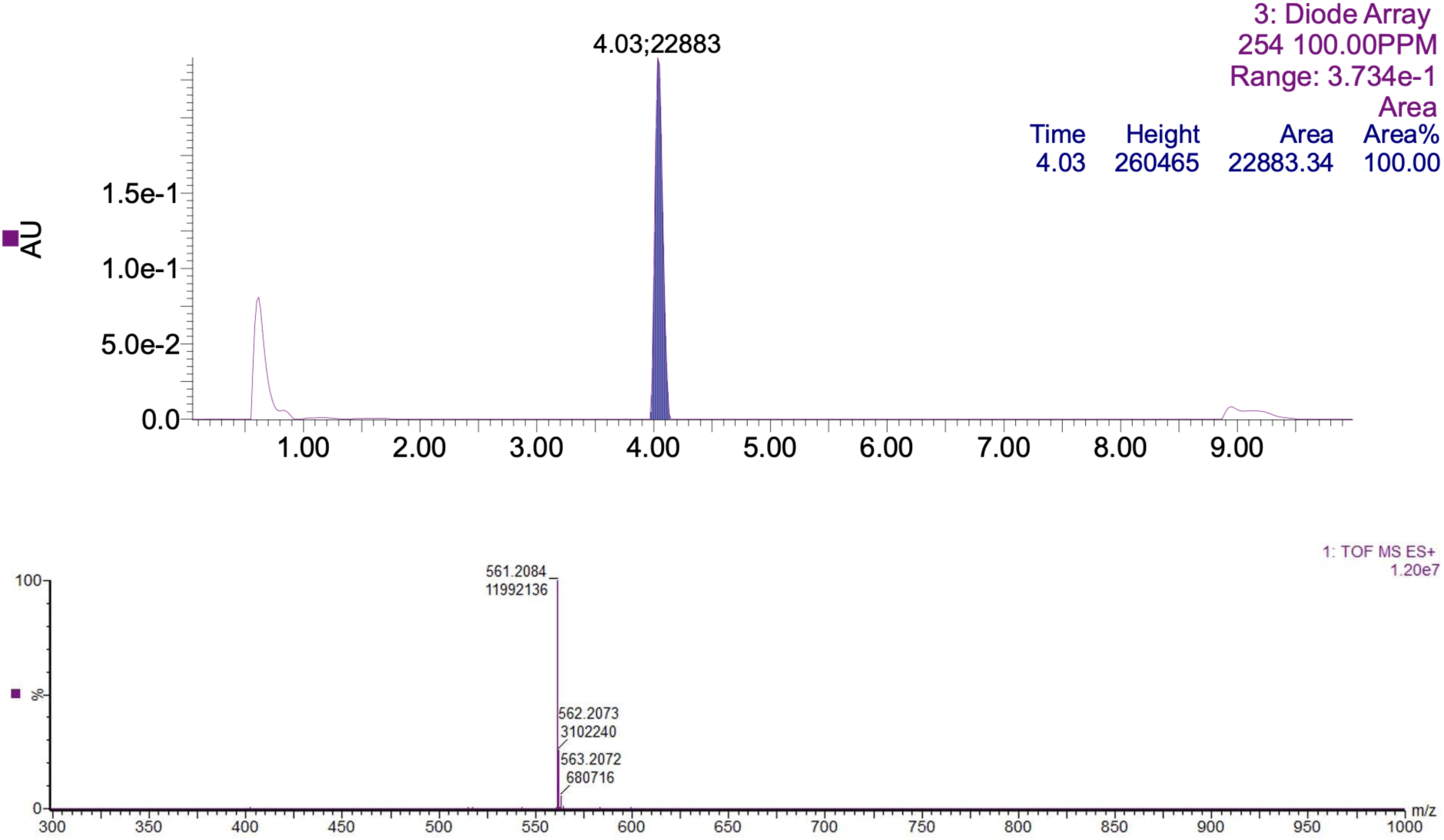

((2S,3S,5R)-3-((1H-1,2,3-Triazol-1-yl)methyl)-3-methyl-4,4-dioxido-7-oxo-4-thia-1-azabicyclo[3.2. **0]heptane-2-carbonyl)-L-phenylalanyl-L-leucine (6)**

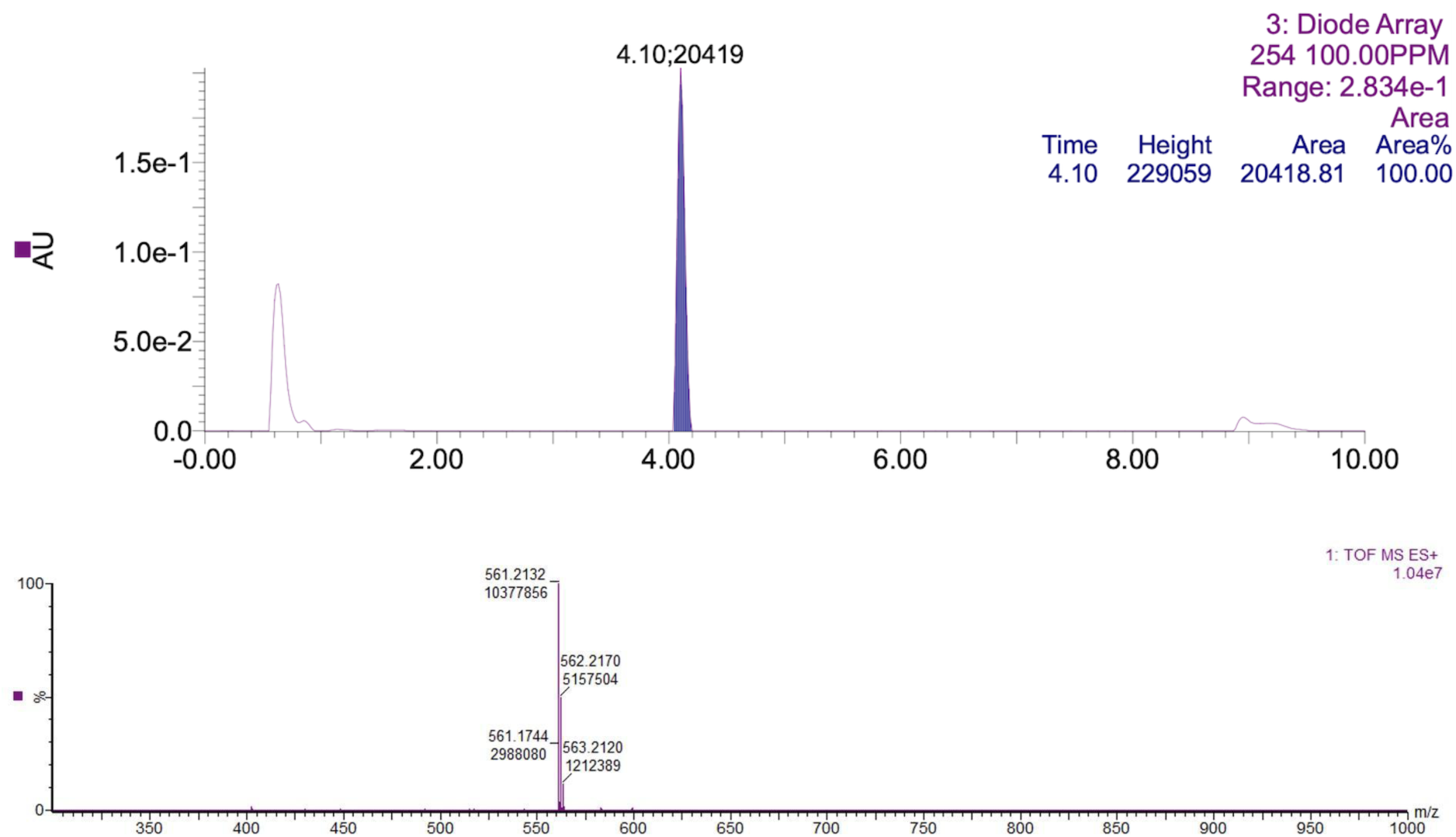

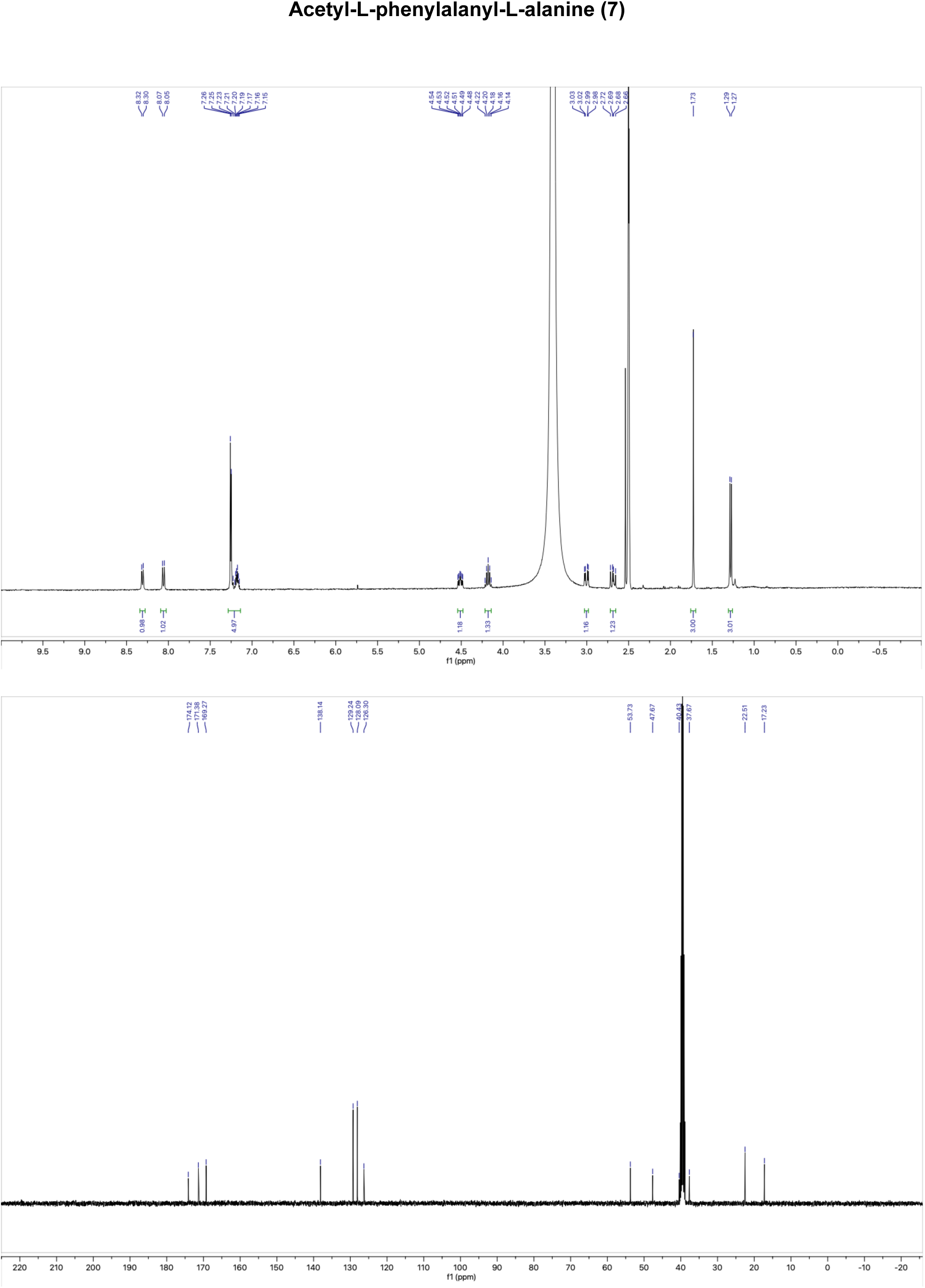

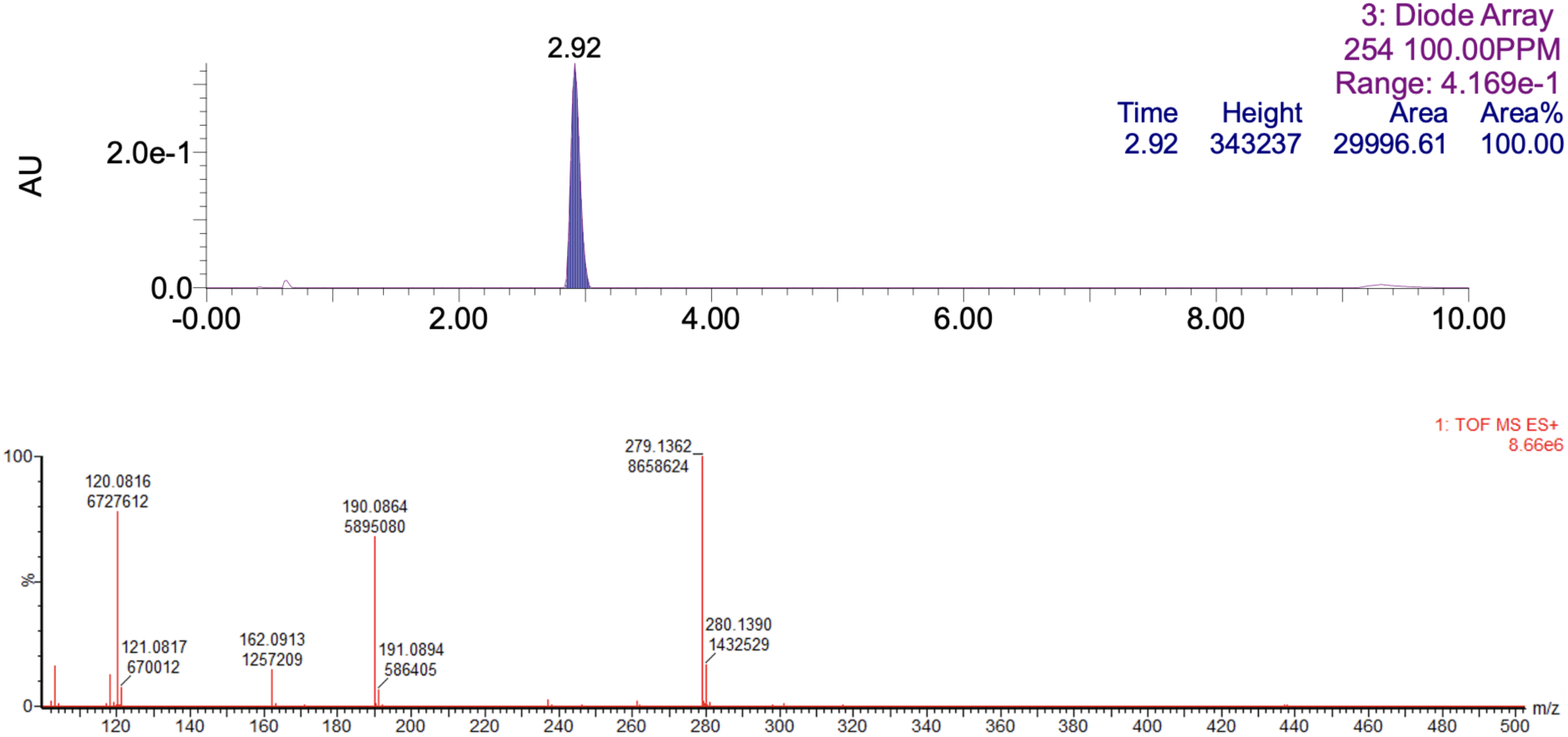

## Notes

### Competing Interest Statement

The authors have declared no competing interest.

